# *Prevotella* are major contributors of sialidases in the human vaginal microbiome

**DOI:** 10.1101/2024.01.09.574895

**Authors:** Paula Pelayo, Fatima A. Hussain, Caroline A. Werlang, Chloe Wu, Benjamin M. Woolston, Katharina Ribbeck, Douglas S. Kwon, Emily P. Balskus

## Abstract

Elevated bacterial sialidase activity in the female genital tract is strongly associated with poor health outcomes including preterm birth and bacterial vaginosis. These negative effects may arise from sialidase-mediated degradation of the protective mucus layer in the cervicovaginal environment. Prior biochemical studies of vaginal bacterial sialidases have focused solely on the bacterial vaginosis-associated organism *Gardnerella vaginalis*. Despite their implications for sexual and reproductive health, sialidases from other vaginal bacteria have not been characterized. Here, we show that vaginal *Prevotella* species produce active sialidases that possess variable activity toward mucin. These sialidases are highly conserved across clades of *Prevotella* from different geographies, hinting at their importance globally. Finally, we find that *Prevotella* sialidases, including mucin-degrading enzymes from *Prevotella timonensis*, are highly prevalent and abundant in human vaginal metagenomes and metatranscriptomes, Together, our results identify *Prevotella* as a critical source of sialidases in the vaginal microbiome, improving our understanding of this detrimental bacterial activity.

**Significance Statement:** Sialidase activity in the vaginal microbiome is increased in bacterial vaginosis and strongly associated with other adverse health outcomes. Sialidase enzymes release sialic acid from host-derived glycans in the vaginal environment, altering their structures and functions. However, biochemical studies of vaginal bacterial sialidases have been limited to one genus, *Gardnerella*. In this work, we identify and characterize multiple active sialidase enzymes in vaginal bacteria of the genus *Prevotella*. We find that *Prevotella* sialidases are more prevalent and abundant in vaginal microbial communities than *Gardnerella* sialidases. Our work highlights *Prevotella* bacteria as an underappreciated source of sialidase activity with important implications for both our understanding of vaginal health and therapeutic development.

## Introduction

The microbial community that inhabits the human vagina (the vaginal microbiome) is important for sexual and reproductive health. The composition of the vaginal microbiome can differ substantially between individuals^1^. While positive health outcomes have been associated with *Lactobacillus-* dominated vaginal microbiomes, more diverse communities containing anaerobic bacteria have been associated with increased risk for preterm birth, bacterial vaginosis (BV)^2^, and HIV acquisition^3^. Despite these strong connections to health, the specific mechanisms by which vaginal bacteria influence the host are poorly understood. An understanding of the vaginal bacterial functions most strongly linked to adverse health outcomes is needed to enhance our understanding of this microbial community and guide the design of vaginal microbiome-targeted therapeutics.

Elevated sialidase activity in vaginal fluid is associated with increased risk of preterm birth^4,5^ and is a common feature of BV^6,7,8^. Sialidases are glycosyl hydrolase enzymes that hydrolyze terminal sialic acids (such as *N*-acetyl-neuraminic acid, Neu5Ac) from glycans present on proteins and lipids (Figure 1A). Multiple, varied sources of sialic acid are found in the female genital tract. Sialic acids are incorporated into the terminal end of mucin glycans, which are prominent components of the mucus layer that covers cervical and vaginal epithelial cells. Immunoglobulins in the cervical mucus (e.g. IgG) are also sialylated, and sialic acids are important for antibody regulation and function^9,10^. Changes in mucus properties are linked to preterm birth risk^11^, suggesting that the activity of sialidases on mucins could have implications for mucus function and host health.

**Figure 1.**
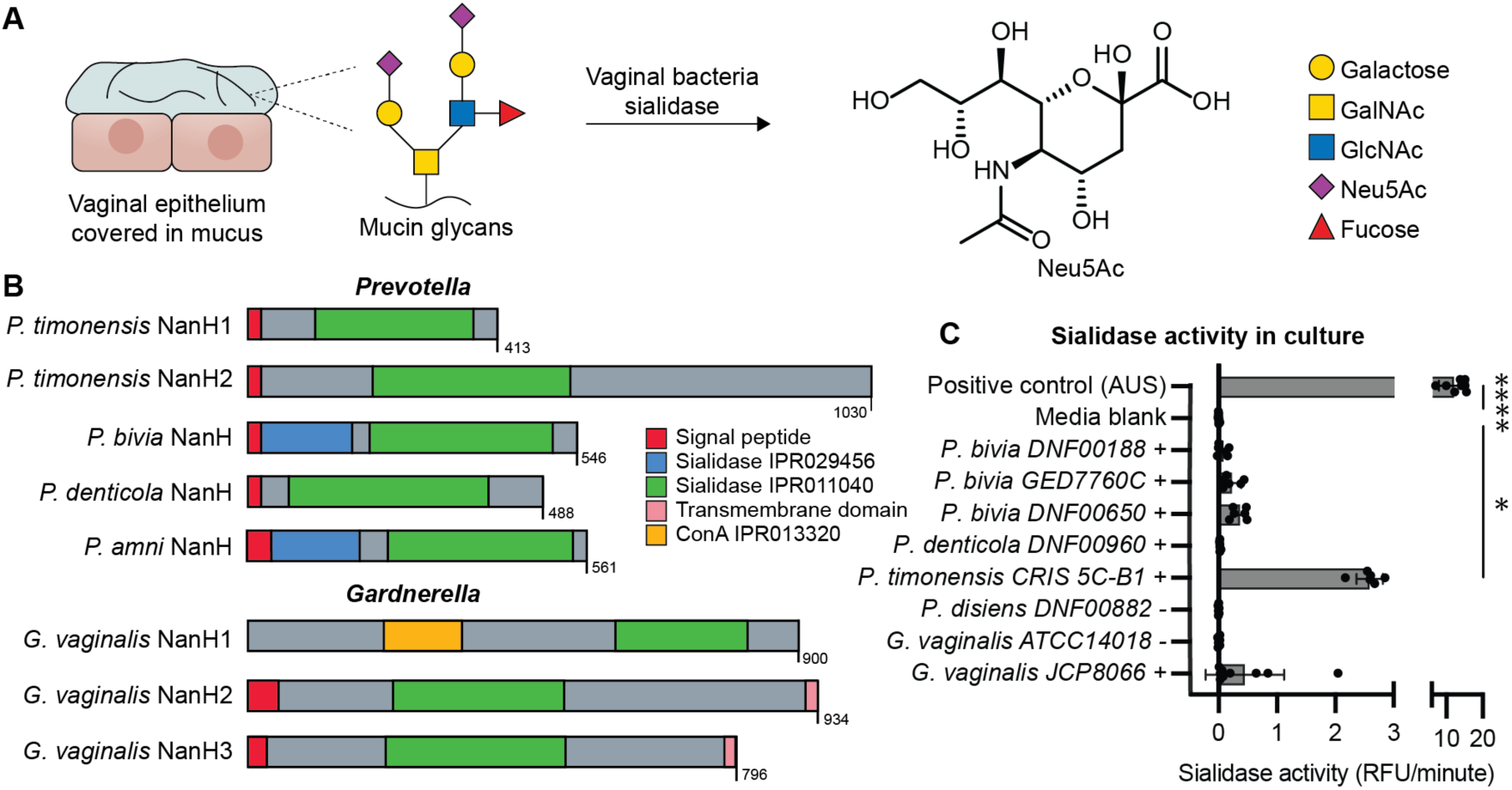
Vaginal *Prevotella* species encode diverse sialidases. (A) Vaginal bacteria sialidase enzymes are hypothesized to remove sialic acids (such as *N*-acetylneuraminic acid, Neu5Ac) from the mucin glycans that comprise the protective mucus layer covering vaginal epithelial cells. (B) *P. timonensis, P. bivia*, *P. denticola*, and *P. amni* encode proteins with predicted sialidase domains. Previously characterized *Gardnerella* sialidases protein domains are also displayed for comparison. Individual colors represent different domains and features: signal peptide (red), sialidase domain IPR011040 (green), and carbohydrate binding domain IPR029456 (blue). The amino acid length is displayed for each bar. (C) *Prevotella* isolates display sialidase activity that correlates with the presence of candidate sialidase genes, indicated by +/–. Positive control represents the full hydrolysis of 4-methylumbelliferyl N-acetyl-α-D-neuraminic acid (4-MU-Neu5Ac) by *Arthrobacter ureafaciens* sialidase (AUS). *G. vaginalis ATCC14018* is indicated as a (–) because it encodes NanH1 but not NanH2 or NanH3. *G. vaginalis JCP8066* encodes NanH1 and NanH3. Data represent the average ± standard deviation (SD) of >5 biological replicates. Significance was assessed using one-way ANOVA followed by multiple comparisons test, ****p<0.0001, *p<0.05. Significance values represent comparison to the media blank.

Sialidase activity in the female genital tract has long been attributed to the anaerobic vaginal bacterium *Gardnerella vaginalis*, which can encode up to three sialidases (NanH1, NanH2, and NanH3)^12^. NanH1 (previously sialidase A), which is highly prevalent in *Gardnerella-*positive vaginal samples^12,13,14^, was originally considered responsible for sialidase activity in this organism. However, recent biochemical characterization of this enzyme showed it has little activity towards the sialidase substrate 4-methylumbelliferyl *N*-acetyl-α-D-neuraminic acid (4-MU-Neu5Ac). NanH1 is also not predicted to be extracellular, making it unlikely to interact with mucin. In contrast, the extracellular enzymes NanH2 and NanH3 display greatly increased activity toward 4-MU-Neu5Ac and can efficiently remove sialic acids from bovine submaxillary mucin^15,12,16^.

Accumulating evidence suggests sialidase activity may be more widespread among vaginal bacteria. For example, several studies have found that *Prevotella bivia*^7^*, Prevotella timonensis* (recently renamed *Hoylesella timonensis*), and *Bacteroides fragilis* strains isolated from vaginal samples possess sialidase activity^6,17,18^. A recent computational study found that *Prevotella* species express a large fraction of predicted sialidases (annotated by the Carbohydrate Active enZymes database, CAZy) in diverse vaginal samples^19^. *Prevotella* strains have been isolated from the upper reproductive tract^20^ and their abundance in the vagina is correlated with preterm birth^21,22^. Persistent sialidase activity in women with recurrent BV has also previously been associated with *P. bivia*^6^. *P. timonensis* isolates can alter the endometrial epithelial barrier and induce mucin expression (MUC3 and MUC4) in a 3D epithelial cell model^18^. *P. timonensis* presence in early pregnancy is also a predictor for higher risk of preterm birth^2^. Despite the intriguing links between sialidase activity, preterm birth, and *Prevotella*, putative sialidases from these bacteria have not been biochemically characterized and their activities remain poorly understood.

Here, we investigate sialidases from vaginal *Prevotella* species as a first step toward understanding their contributions to sialidase activity in the female genital tract. We initially biochemically characterize sialidases from three *Prevotella* species *in vitro*, observing unexpected differences in activity toward mucin substrates, including the human mucin MUC5B. We use comparative genomics to demonstrate that *Prevotella* sialidases are widely distributed and conserved across vaginal isolates from the United States and South Africa. Finally, through analysis of human vaginal metagenomes and metatranscriptomes, we discover that *P. timonensis* sialidases are encoded and expressed at a higher prevalence and abundance than sialidases from other vaginal bacteria, including *G. vaginalis*. These findings reveal *Prevotella* bacteria as important, underappreciated contributors to sialidase activity in the human vaginal microbiome and highlight a need to understand the biological roles of these enzymes in the vaginal environment.

## Results

### Vaginal *Prevotella* species encode diverse sialidases

To identify candidate sialidase genes in vaginal *Prevotella*, we initially analyzed seven genomes of from vaginal isolates of *P. bivia, P. amnii, P. denticola, P. disiens*, and *P. timonensis.* Sequences of biochemically characterized sialidases from *G. vaginalis* (NanH2 and NanH3) were used as query. We found five genes encoding candidate sialidases in four *Prevotella* species, *P. bivia* (*PbnanH*), *P. amnii* (*PananH*), *P. denticola* (*PdnanH*), and *P. timonensis* (*PtnanH1* and *PtnanH2*) (Figure S1). All five proteins contain the catalytic active site residues characteristic of the glycoside hydrolase 33 (GH33) family of sialidases (Figure S2, Table S5). All sequences also contain a predicted signal peptide, indicating they are likely extracellular, and a sialidase catalytic domain (IPR011040) (Figure 1B) with the predicted β-propeller fold characteristic of these enzymes (Figure S3-S5). *PbnanH* and *PananH* also contain a predicted carbohydrate-binding domain, which is typically found in sialidases from commensal gut *Bacteroides*^23^. We found that *P. timonensis* encodes two sialidases of different lengths (*Pt*NanH1 413 amino acids; *Pt*NanH2 1,030 amino acids) and AlphaFold^24^ structure prediction indicates PtNanH2 may contain four additional domains of unknown function (Figure S5B).

We next cultured available vaginal *Prevotella* isolates and tested for sialidase activity using the fluorescent substrate 4-MU-Neu5Ac. Notably, of all the bacterial cultures tested, *P. timonensis* consistently had the highest sialidase activity (Figure 1C). Sialidase activity in *G. vaginalis* JCP8066, which encodes NanH3, was detectable but variable across different experiments. Other *Prevotella* strains encoding sialidases had low but detectable activity. No activity was observed for a *P. disiens* isolate lacking a sialidase homolog. Together, this work further confirmed the presence of sialidase activity in vaginal *Prevotella* strains and identified candidate sialidase enzymes for biochemical studies.

### *Prevotella* sialidases are active and susceptible to inhibition

To determine if the predicted *Prevotella* sialidase genes encoded active sialidase enzymes, we expressed the putative sialidases from *P. timonensis, P. bivia*, and *P. denticola* in *E. coli* and purified them for *in vitro* biochemical characterization (Figure S6). We found all purified *Prevotella* sialidases were active toward 4-MU-Neu5Ac (Figure 2A). The kinetics of vaginal bacterial sialidases have not yet been examined; we therefore determined the Michaelis–Menten kinetic parameters of the *Prevotella* sialidases and *Gv*NanH3. We found their turnover rates *k*_cat_ (110.46– 149.33 s^-1^) and catalytic efficiencies k_cat_ / K_m_ (0.51–1.83 x 10^6^ s^-1^ M^-1^) (Table S6) comparable to those of other previously characterized bacterial sialidases (Table S3).

**Figure 2.**
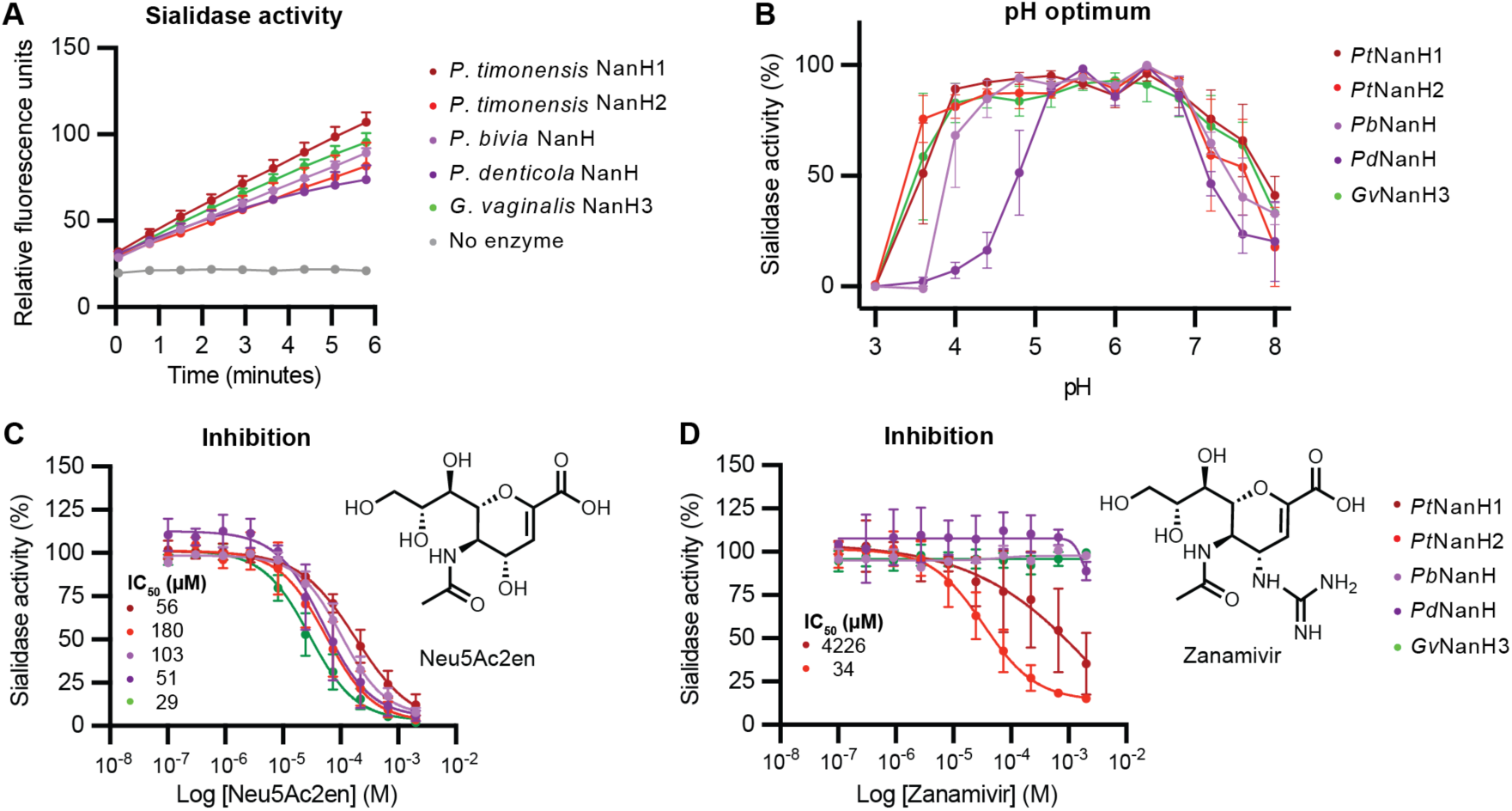
*Prevotella* sialidases are active at varying pH levels and are inhibited by small molecules. (A) Purified sialidase enzymes hydrolyze 4-MU-Neu5Ac, indicating they possess sialidase activity. For each time point, 2.5 nM enzyme was incubated with 200 µM 4-MU-Neu5Ac in sodium acetate buffer, pH 5.5 at 37 °C. Data represent the average + standard error (SEM) of 3 independent experiments. (B) Sialidase activity at varying pH. 200 µM 4-MU-Neu5Ac was prepared in 0.1 M citric acid / 0.2 M phosphate buffer at pH values ranging from 3–8. Data represent the average ± SEM of 3 independent experiments. Inhibitory activity of (C) Neu5Ac2en and (D) Zanamivir towards *Prevotella* and *Gardnerella* sialidases (IC_50_ values can be found in Table S8). Purified sialidases were pre-incubated with inhibitor for 15 minutes before adding 4-MU-Neu5Ac to determine activity. Data represent the average ± SEM of 3 independent experiments.

Healthy *Lactobacillus*-dominated vaginal communities have a pH <4.5^1^, while *Prevotella* species are typically found in more diverse vaginal communities which have a pH >4.5. We therefore examined the pH dependence of the *Prevotella* sialidases and *Gv*NanH3. Measuring the activity of the *Prevotella* sialidases toward 4-MU-Neu5Ac over a pH range of 3.0 to 8.0 revealed that all sialidases had maximal activity above pH 4.5 (Figure 2B). Interestingly, among the tested enzymes *Pt*NanH1, *Pt*NanH2, and *Gv*NanH3 displayed activity below pH 4 and, in contrast, *Pd*NanH was only active above pH 4.8.

To further confirm the functional assignment of these enzymes, we examined their susceptibility to various commercially available sialidase inhibitors. All enzymes were inhibited by the well characterized, broad spectrum sialidase inhibitor, Neu5ac2en (Figure 2C, Table S4)^25^. Unexpectedly, the viral sialidase inhibitor Zanamivir (Relenza) was effective towards *P. timonensis* sialidases *Pt*NanH1 and *Pt*NanH2 but no other enzymes tested (Figure 2D). Inhibition of sialidase activity by Neu5Ac2en and Zanamivir was also observed in *Prevotella* cultures (Figure S8). Overall, these results further support the characterization of these enzymes as sialidases, with the variable activity of Zanamivir suggesting potential differences in structure and function between the *P. timonensis* sialidases and those from other *Prevotella* species.

### *Prevotella* sialidases have variable activity towards mucin glycoproteins

Sialic acids are incorporated into a variety of potential substrates, including mucin glycans, via α2-3’ and α2-6’ linkages. We sought to determine if *Prevotella* sialidases had differences in linkage and substrate preferences by testing their activity toward a panel of six sialic acid-containing substrates (Figure 3A). We incubated purified sialidases with α2-3’-sialyllactose (3’SL), α2-6’-sialyllactose (6’SL), bovine submaxillary mucin (BSM), human salivary MUC5B, immunoglobulin A (IgA), and immunoglobulin G (IgG), and quantified Neu5Ac released by each enzyme in an endpoint liquid chromatography–mass spectrometry (LC–MS) assay. We found that all sialidases released sialic acid from 3’SL and 6’SL substrates, indicating these sialidases do not have a strong preference for linkage type (Figure 3A). All sialidases tested were able to release sialic acid from IgA and IgG, the predominant immunoglobulin in cervical mucus (Figure 3A). Unexpectedly, the *Prevotella* sialidases differed in their activity toward mucin substrates. While *Pt*NanH2 and *Gv*NanH3 efficiently removed sialic acid from both BSM and human MUC5B, *Pt*NanH1, *Pb*NanH and *Pd*NanH had reduced activity (30-40%) toward BSM and little to no activity toward MUC5B (Figure 3B and 3C). The variable activity of vaginal bacterial sialidases toward the human mucin MUC5B is unexpected and striking given the potential links between mucin degradation and negative health outcomes.

**Figure 3.**
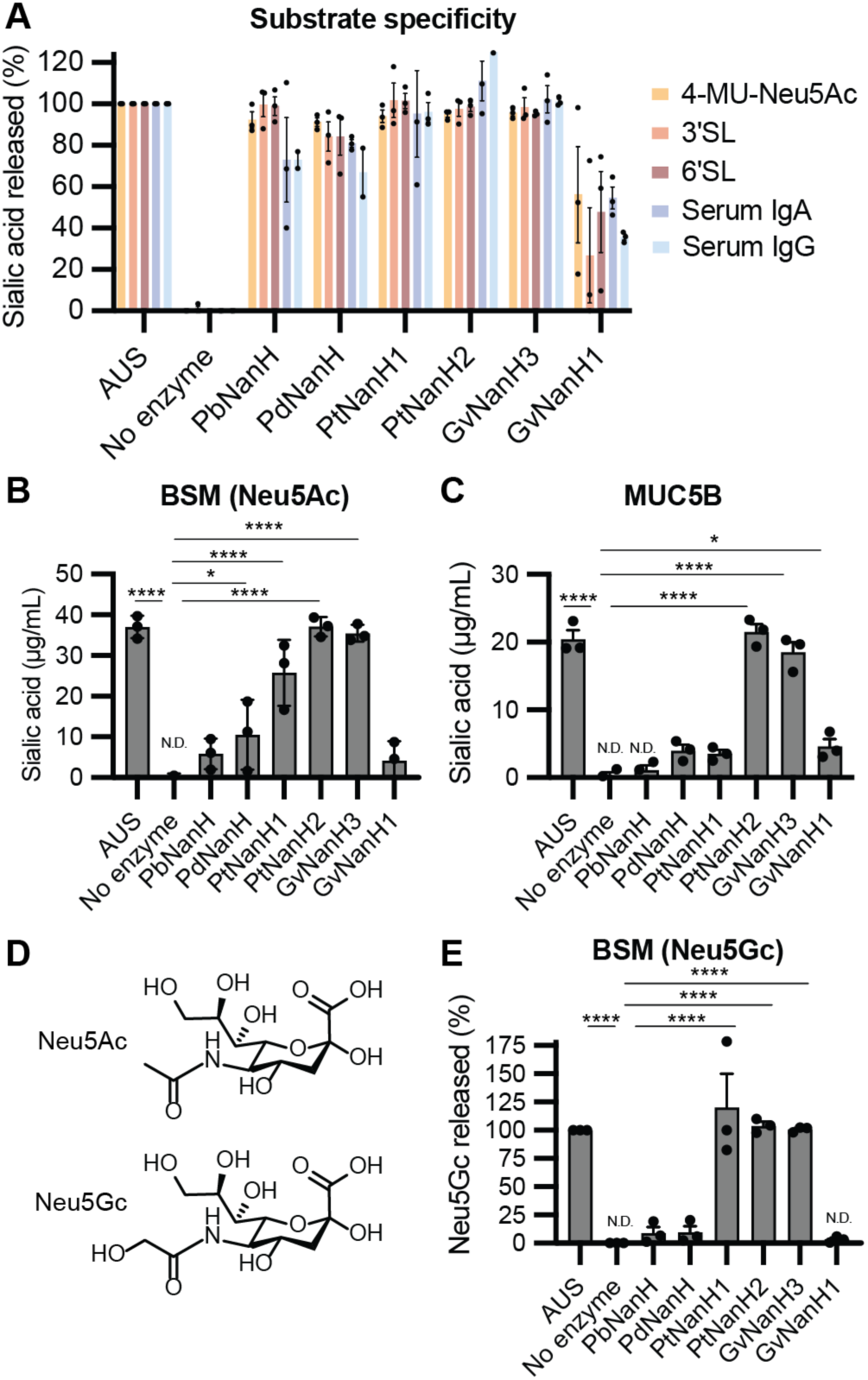
*Prevotella* sialidases have varied substrate specificities. (A) Quantification of sialidase activity towards different substrates: 4-MU-Neu5Ac, α2-3’siallylactose (3’SL) and α2-6’siallylactose (6’SL), human serum immunoglobulin G (IgG), and human serum immunoglobulin A (IgA). Purified sialidase enzyme (20 nM–100 nM) was incubated with each substrate in 20 mM sodium acetate buffer pH 5.5 for 2 hours at 37 °C. *Arthobacter ureafaciens* sialidase (AUS) was used a positive control to release the total sialic acid from each substrate^43^. Sialidase activity towards (B) bovine submaxillary mucin (BSM) and (C) purified human salivary MUC5B. (D) Structures of *N*-acetylneuraminic acid (Neu5Ac) and *N*-glycolylneuraminic acid (Neu5Gc). Data represent the average ± SEM of 3 independent experiments. N.D. represents values were not detected below the limit of detection. (E) Sialidase activity towards Neu5Gc from BSM. Data represent the average ± SEM of 3 independent experiments. Significance was assessed using one-way ANOVA followed by multiple comparisons test, ****p<0.0001, *p<0.05. Significance values represent comparison to the no enzyme control.

We next sought to explore whether *Prevotella* sialidases could release other forms of sialic acid from BSM, which contains diverse sialic acid structures including Neu5Ac and Neu5Gc, Neu5,7Ac_2_, and Neu5,9Ac_2_. *Pb*NanH resembles Sialidase26 (66% amino acid ID), recently characterized from the human gut metagenomes^26^, which has preferential activity towards *N*-glycolylneuraminic acid (Neu5Gc) (Figure 3D). While the presence of Neu5Gc in cervicovaginal mucins is unclear, this prompted us to assess whether these sialidases could release Neu5Gc from BSM. Consistent with our previous assays examining activity toward BSM, *Pb*NanH and *Pd*NanH did not release Neu5Gc from BSM (Figure 3E). Among the sialidases observed previously to release Neu5Ac from BSM, only *Pt*NanH2 and *Gv*NanH3 also released Neu5Gc from this substrate, further indicating these enzymes are more promiscuous than the other sialidases (Figure 3E). Together, the results of this substrate survey demonstrate *Prevotella* sialidases can release sialic acid from substrates resembling those found in the cervicovaginal environment. Notably, the *P. timonensis* enzymes possess similar reactivity to the previously reported active *Gardnerella* sialidase *Gv*NanH3^12^. Importantly, the wide variation in sialidase substrate specificities we observed could not have been anticipated based on their annotation, further underscoring the importance of *in vitro* biochemical studies in understanding enzymes from the human microbiome.

### Sialidases are conserved in *Prevotella* isolates from different geographies

Having characterized the activities of the *Prevotella* sialidases, we next we sought to determine their prevalence across diverse strains of *Prevotella* isolated from different geographies and compare their distribution to that of other characterized and predicted vaginal bacterial sialidases. Employing an HMM-based approach using the sialidases characterized in this work and other GH33 sialidases from the CAZy database, we searched whole genomes from over 1,000 bacterial strains isolated from different cohorts in the United States and South Africa (Figure S9). We observed that *P. timonensis* and *P. bivia* sialidases are highly conserved within isolates of these species, with all 53 *P. bivia* isolates encoding the same PbnanH sialidase (>98% amino acid ID) regardless of the geography of origin. Similarly, all 21 *P. timonensis* isolates encode PtnanH2 (>93% amino acid ID) and 19/21 encode PtnanH1 (>94% amino acid ID) (Figure 4A). Additionally, we found close relatives of *P. timonensis* also encode distantly related sialidases. A vaginal *P. buccalis* isolate encodes a sialidase with 61% amino acid ID to *Pt*NanH1 but lacks a signal peptide. *P. colorans FRESH097* and *P. sp16 C0026C3* encode a putative sialidase distinct from the ones characterized here with only 9% and 23% amino acid ID to *Pt*NanH1, respectively, and both contain signal peptides (Figure S11). Interestingly, we did not detect any genes encoding sialidases in clades of *P. disiens*, *P. intermedia, P. ihumii,* or *P. corpo*ris, and we did not find additional examples of vaginal *P. denticola* genomes. Inspecting the gene neighborhoods of these *Prevotella* sialidases revealed they are usually found near genes encoding other glycoside hydrolase enzymes and are not usually co-localized with sialic acid metabolism genes, except for *P. timonensis nanH2* which is near *nanE* (UDP-N-acetylglucosamine 2-epimerase) (Figure 4B).

**Figure 4.**
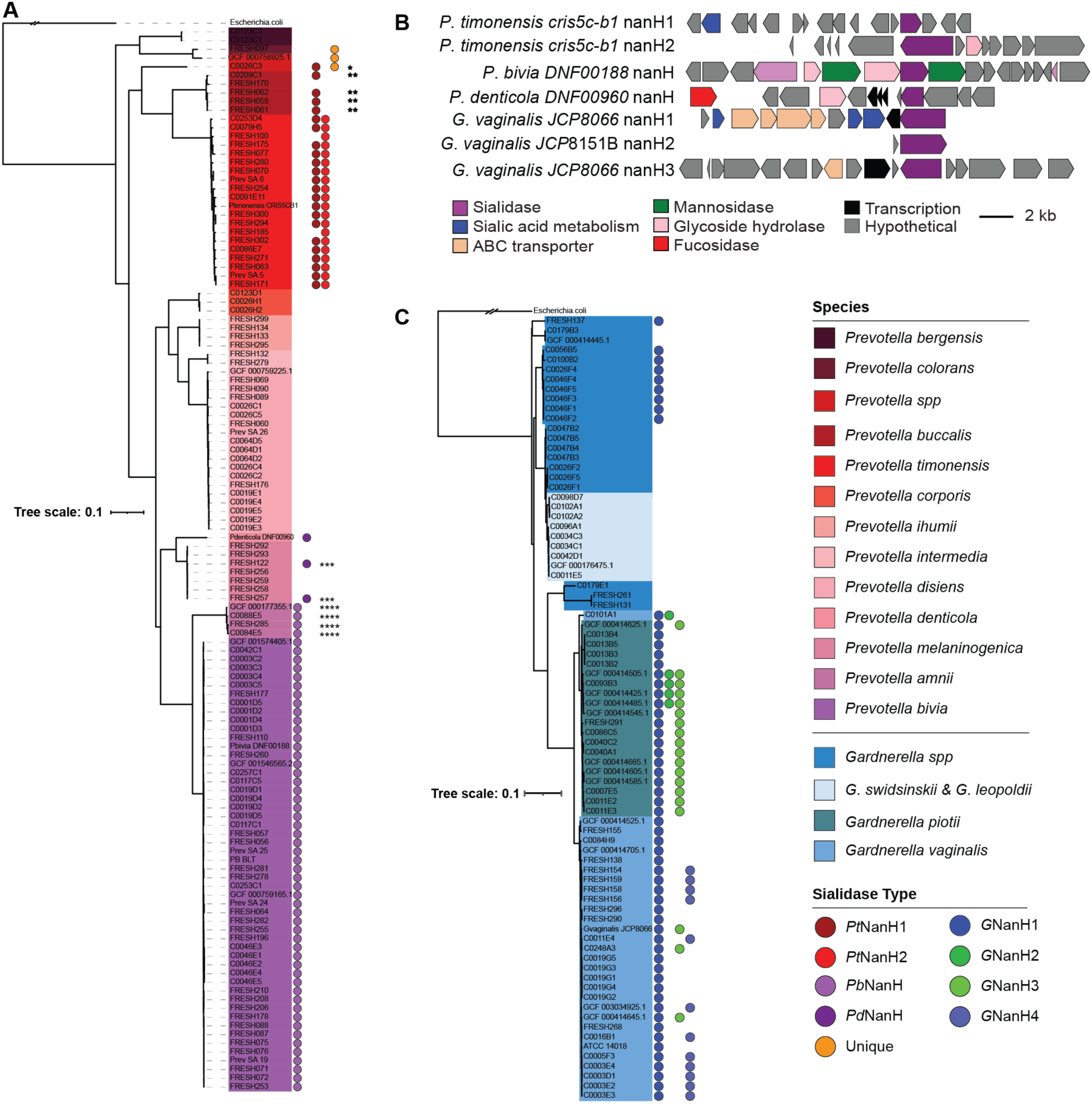
Sialidases are widely distributed across *Prevotella* and *Gardnerella* isolates obtained from the United States and South African vaginal samples. Phylogenetic trees of vaginal isolates from the Vaginal Microbiome Research Consortium (N. American) and FRESH (S. African) studies. The sialidase genes present in individual (A) *Prevotella* and (B) *Gardnerella* isolate genomes are shown as filled circles. The phylogeny is based on 49 concatenated ribosomal proteins and serves as a proxy for the core genome. The scale bar indicates nucleotides substitutions per site. Each genome was searched using the sialidase protein alignment with HMMER (version 3.1b2). (C) Genome neighborhoods of *Prevotella* and *Gardnerella* sialidases.*=56% amino acid ID to *Pt*NanH1, **=61% AA ID to *Pt*NanH1, ***=78% amino acid ID to *Pd*NanH, ****=70% amino acid ID to *Pb*NanH. The orange filled circle indicates a sialidase that is unique from the *Prevotella* sialidases characterized here.

*Gardnerella* isolates varied in their sialidase content based on clade, as reported previously^27^. *G. swidsinskii* and *G. leopoldi* did not encode any sialidases, while most *G. vaginalis* and *G. piotii* encode *GvnanH1* (61/73). Notably, the active sialidases *GvnanH2* (5/73) and *GvnanH3 (*20/73) are much less prevalent than *GvnanH1* and are unevenly distributed across strains of individual species (Figure 4C). Interestingly, this analysis revealed variants of *GvnanH1* exist among different *Gardnerella* clades. For example, a *G. vaginalis* variant of NanH1 found in isolates from the United States is 82 % amino acid ID to *G. vaginalis* NanH1 found in strains from South Africa. *G. vaginalis JCP8066* NanH3 is homologous to *G. piotii* NanH3 (92% amino acid ID). We also identified an additional candidate sialidase, called NanH4, in 12 *G. vaginalis* isolates; however, NanH4 may be inactive since it is found in *G. vaginalis ATCC 49145,* which has not demonstrated any sialidase activity^12^.

We also found putative GH33 sialidases among diverse vaginal isolates from the genus *Bacteroidales, Bacteroides, Bifidobacterium, Corynebacterium (Actinomycetia*), and *Streptococcus* (Figure S9-10). *Streptococcus agalactiae* encodes a sialidase that is >98% ID to Group B *Streptococcus* NonA^28^, an inactive homolog of the *S. pneumoniae* NanA sialidase. There was no sialidase present in its sister clade *S. anginosus*. Overall, this genome survey increases knowledge of vaginal bacterial sialidases by broadening the search to a wide variety of species and geographies. The substantial variability we observe in sialidase presence across certain vaginal bacterial species highlights a need to directly identify these enzymes in vaginal microbiomes rather than infer their presence from phylogenetic information.

### *P. timonensis* sialidases are prominent in vaginal microbiomes

We next identified the biochemically characterized *Prevotella* and *Gardnerella* sialidases, as well as the additional putative sialidases from our genome survey, in human vaginal microbiomes. To quantify the differential abundance and prevalence of sialidases, we performed translated nucleotide sequence searches in paired metagenomes (MG) and metatranscriptomes (MT) from vaginal samples sequenced by France *et al*. ^19^ (Figure 5A). These samples were collected from 39 reproductive aged, non-pregnant women at up to 5 timepoints over the span of 10 weeks. We analyzed all the samples individually and paired the MG and MT samples (n=176) with the associated subject metadata to examine association of sialidase expression with community state type (CST), including *L. crispatus*-dominated (CSTI), *L. gasseri*-dominated (CSTII), *L. iners*-dominated (CSTIII), diverse anaerobic (CSTIV), and *L. jensenii*-dominated (CSTV) communities. We expected to find higher levels of sialidases in CSTIV communities compared to other CSTs since sialidase activity is detected in subjects with BV and is typically absent from *Lactobacillus*-dominated samples^6,29^. Unexpectedly, sialidases were detected in a high percentage of all CSTs (69–100% of MGs and 56–90% of MTs). However, we found that CSTIV MGs encode a greater abundance of sialidases compared to all other CSTs (Figure 5A). We also found that CSTIV MT samples have significantly higher sialidase expression than MTs from CSTI and CSTIII samples (Figure 5B).

**Figure 5.**
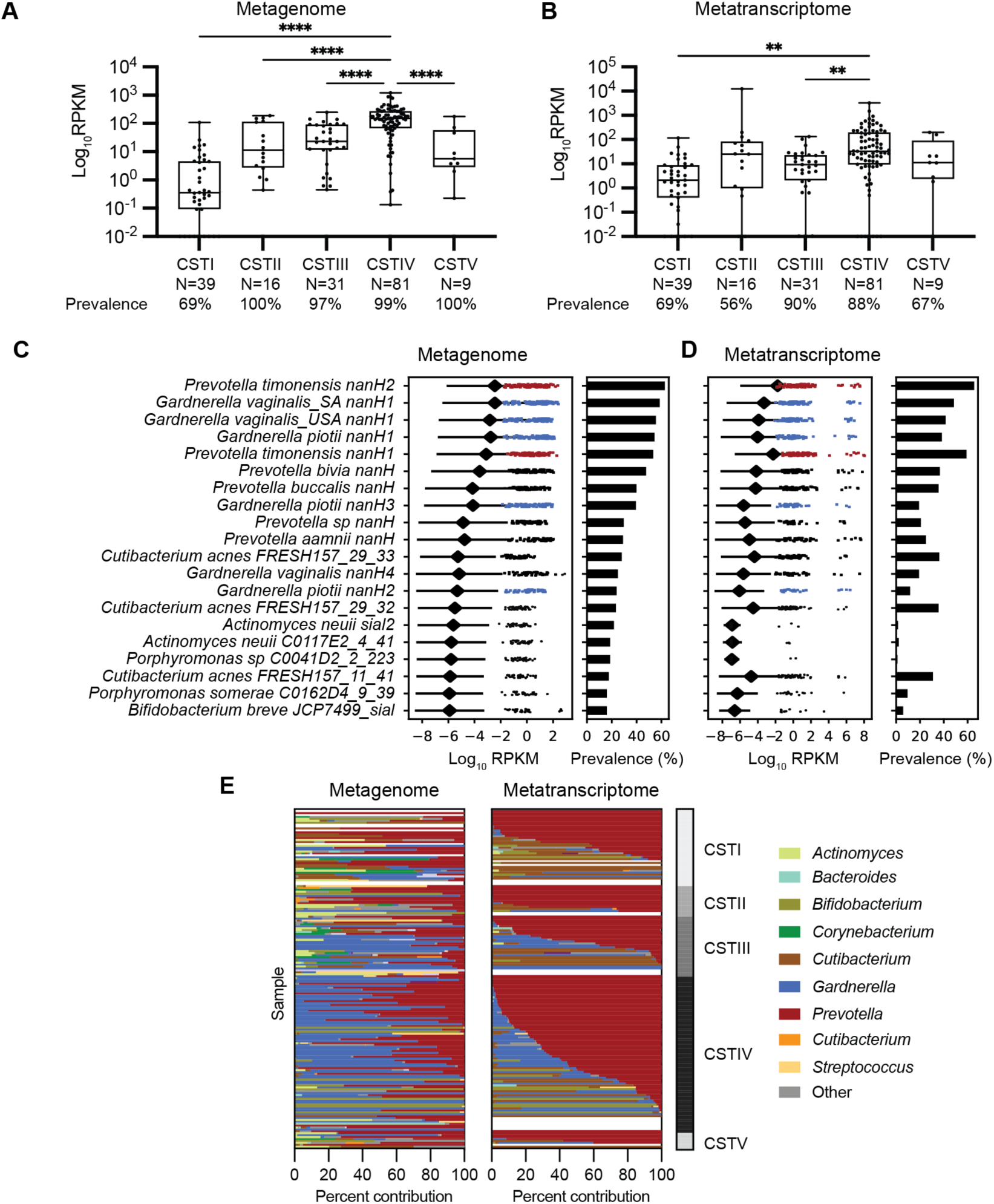
*P. timonensis* sialidases are prevalent across vaginal community state types. (A) Total sialidase abundance and prevalence in paired metagenome and (B) metatranscriptomes samples (n=176). Abundance was determined by Diamond blastX and displayed as reads per kilobase million (RPKM). Significance was assessed by a one-way ANOVA followed by a Brown-Forsythe and Welch test, **p<0.0001, **p<0.01. (C) The abundance and prevalence of specific sialidase genes in all available samples (MG; n=193, MT; n=188). Figure displays sialidases with >15 % prevalence in MG samples. The diamond indicates the average sialidase expression across all samples. (E) Contribution of sialidase abundance within individual vaginal samples across metagenomes and metatranscriptomes. Paired metagenomes (MG) and metatranscriptomes (MT) were used to investigate the relative contribution of several vaginal sialidases. Stacked bar graphs are aligned to pair the corresponding MG and MT sample.

The prevalence of individual sialidase genes varied across all vaginal MGs: *G. vaginalis* sialidases were detected in 23%–58% of samples, *P. bivia* (*PbnanH* 47% of samples), *P. amni* (*PananH* 29% of samples), and *P. timonensis* sialidases (*PtnanH1* 53% and *PtnanH2* 62% of samples) were identified at similar frequencies, while the sialidase from *P. denticola* was not detected (0%), likely because this organism is rarely found in the vaginal microbiome (Figure 5C). In vaginal MTs, *P. timonensis* sialidase transcripts were the most prevalent (*PtnanH1* 59% and *PtnanH2* 65% of samples) and abundant, followed by *G. vaginalis* (*GvnanH1* 38–48%, *GvnanH2* 11%, *GvnanH3* 19% of samples), *P. amni* (*PananH* 25% of samples) and *P. bivia* (*PbnanH* 36% of samples) (Figure 5D). Interestingly, *G. vaginalis nanH1* variants were more prevalent than the active *Gardnerella* sialidases (*G. pioti nanH2* and *nanH3*). Sialidases from additional taxa identified in Figure 4 have low prevalence in MGs (<15% of samples) and had minimal to no presence in the MTs.

The variable prevalence of different sialidases across samples raised the question of their contributions to overall sialidase expression within individual samples. To determine which sialidases are expressed in individual vaginal microbiomes, we computed the percent contribution of all sialidases within paired MG and MT samples (Figure 5E). We found that, although *Gardnerella* sialidases are present in MGs, *P. timonensis* sialidases and other *Prevotella* sialidases contribute a higher percentage of reads in MTs across samples from all CSTs (Figure 5E, Figure S12A). *P. timonensis* sialidases were also more prevalent than all *G. vaginalis* sialidases across all CSTs (Figure S12). Together these findings show that while both *Prevotella* and *Gardnerella* sialidase genes and transcripts are detected in vaginal samples, those from *Prevotella* are most prevalent and abundant across all CSTs. Importantly, this pattern is not simply a reflection of vaginal microbiome community composition, as previous analyses of these datasets revealed a higher abundance of *Gardnerella*^19^. Overall, our analyses indicate that *Prevotella* bacteria, particularly *P. timonensis*, are a predominant source of sialidase activity within the vaginal microbiome and should be considered in developing therapeutic interventions.

## Discussion

Sialidase activity in the vaginal microbiome has been strongly linked to negative health outcomes, including BV and preterm birth. Thought to significantly alter the vaginal ecosystem, this metabolic activity may facilitate mucus degradation, release carbon sources for vaginal bacteria, and promote bacterial binding to mucins and host cell glycans by revealing cryptic binding sites^30^. Indeed, a recent study found that human vaginal epithelial surface has a diminished glycocalyx during BV and this phenotype can be recapitulated in cell culture with recombinant *Gardnerella* sialidases, further implicating these enzymes in detrimental host phenotypes. Prior to our work, knowledge of sialidase enzymes in vaginal bacteria was largely limited to *Gardnerella*. *Gardnerella* encode three sialidases, NanH1 (which is poorly active) as well as NanH2 and NanH3, which display sialidase activity and are predicted to localize extracellularly. Other vaginal bacteria were reported to possess sialidase activity in culture, including *P. bivia* and *P. timonensis*^6,18^. Here, we combine bioinformatic analyses and biochemical experiments to identify *Prevotella* sialidase enzymes and assess their distribution across vaginal isolates and vaginal microbiomes.

Though enzyme encoding genes can be annotated in bacterial genomes, important aspects of their activity cannot be readily predicted from sequence alone, highlighting the importance of biochemical characterization. Indeed, our *in vitro* studies of vaginal sialidases revealed differences in pH activity range, activity towards mucin substrates, and susceptibility to inhibition. Our pH analysis reveals *P. timonensis Pt*NanH1 and *Pt*NanH2 are active at pH values <4.5 and >4.5, while the other bacterial sialidases tested only displayed activity at pH >4.5. This finding suggests that *P. timonensis* sialidases may be active in health-associated *Lactobacillus*-dominated communities. We discovered that only a subset of *Prevotella* sialidases, including enzymes from *P. timonensis*, accept mucin substrates perhaps indicating a higher potential to alter the structure and function of these important host glycans. While the broad-spectrum inhibitor Neu5Ac2en inhibited all sialidases, the viral sialidase inhibitor Zanamivir specifically inhibited both *P. timonensis* sialidases. Though we did not observe inhibition of GvNanH3 by Zanamivir in our experimental setup, it is possible that this enzyme can be partially inhibited as was previously demonstrated^31^. In an accompanying paper^32^, Segui Perez et al., demonstrate that removal of sialic acids from vaginal and epithelial cell surface glycans by *P. timonensis* can be prevented by treatment with Neu5Ac2en and Zanamivir. Sialidase inhibitors are therefore useful tools to understand the effects of sialidase activity in the vaginal environment and could be further explored therapeutically,

We found *Prevotella* sialidases are more widely conserved across individual strains compared to the *Gardnerella* sialidases. Previous studies identified sialidase activity in most *P. bivia* isolates^27^, but few studies have examined *P. timonensis*. Using a diverse isolate genome collection also allowed us to find additional putative sialidases from closely related *Prevotella* species. We predict the newly identified *Gardnerella* sialidase *Gv*NanH4 is likely inactive, similar to *Gv*NanH1, since no strains encoding this gene are known to have sialidase activity. *Gv*NanH4 and *Gv*NanH1 have predicted Ig/Lectin-like domains that can also be found in *Actinomyces* sialidases, which have not been biochemically characterized. It is possible that *Gv*NanH4 acts intracellularly on an as-yet-uncharacterized substrate given its lack of a signal peptide, or is inactive, similar to the Group B Streptococcus sialidase NonA^28^. Future work should examine sialidase activity towards additional substrates such as human vaginal epithelial cell surface glycans^33^, glycan arrays^34^, or in more complex or natural settings^35,18^.

Our analyses of vaginal MGs and MTs demonstrates most of the sialidase expression comes from *Prevotella* and *Gardnerella* while putative sialidases from other vaginal bacteria are rarely expressed. Among the *Gardnerella* sialidases, *GvnanH1* was the most prevalent in MTs. This is perhaps surprising considering experimental evidence shows GvNanH1 has extremely low activity toward sialylated substrates^12^. *GvnanH1* may be more prevalent in MTs because it is more frequently found in *Gardnerella* genomes compared to *GvnanH2* and *GvnanH3*^36^; however, it may also participate in an uncharacterized metabolic process. Notably, *Gardnerella’s* active sialidases, *GvnanH2* and *GvnanH3,* were less prevalent than *PtnanH1* and *PtnanH2* in this dataset, suggesting they may have less of an impact on host glycan metabolism. Interestingly, in the accompanying paper^32^, Segui Perez et al. demonstrated *P. timonensis* has much higher cell-bound and secreted sialidase activity in culture and is more active against cell surface glycans compared to *G. vaginalis* and *P. bivia*. Together, these data suggest that, in addition to *Gardnerella*, *P. timonensis* make a substantial contribution to vaginal sialidase activity, calling into question previous assumptions about its origins.

Sialidases activity is considered a hallmark of BV^8^ and diverse CSTIV communities. Our data suggest sialidases have the highest prevalence in CSTIV samples, however, we observed sialidase expression in samples from other CSTs as well (Figure S12). Although rare, several studies have detected sialidase activity in non-BV samples^6,37,38^. We also noticed samples can vary in their sialidase expression profile, with some expressing three to four sialidases, while others express a single sialidase. This hints at the relevance of multiple, diverse sialidases in the vaginal environment, potentially creating a division of labor or even the potential for public good exploiters as seen in other ecosystems^39^. Other studies have noted while the relative abundance of *Prevotella* species in vaginal MGs may be low^1^, their genes can be highly expressed in MT data^19^. In particular, the presence of *P. timonensis* sialidase transcripts in MTs from certain *Lactobacillus*-dominated communities suggests the possibility that these enzymes could contribute to community destabilization and transition to other CSTs.

Together, these bioinformatic analyses provide a more complete understanding of the origins of sialidase activity in the vaginal microbiome and highlight the importance of integrating bioinformatic analysis with detailed biochemical studies. Additional analyses of vaginal bacterial sialidase genes and transcripts in other clinical cohorts are needed, including pregnancy cohorts that could reveal links between specific sialidases and birth outcomes.

In summary, we demonstrate that *Prevotella* possess sialidase enzymes that likely play an important, currently underappreciated role in the vaginal microbiome, with *P. timonensis* standing out as a prominent source of sialidase enzymes and a major contributor of sialidase transcripts in vaginal communities. In the accompanying manuscript, we also demonstrate that *P. timonensis* can efficiently adhere to the vaginal epithelium and that its sialidases and fucosidases are highly effective at removing glycans from the vaginal epithelial surface^32^. While *Gardnerella* has been previously considered primarily responsible for sialidase activity in this environment, our work indicates that *Prevotella* species should also be included in future studies and considered in developing therapeutics targeting the vaginal microbiome. These discoveries also highlight the need for further investigation into the biological roles of vaginal bacterial sialidases and their contributions to negative health outcomes.

## Materials and Methods

### Plasmid Construction

Full length sialidase genes were amplified (Q5 polymerase, New England Biolabs, M0492S) from purified genomic DNA (Qiagen, 12224-50) from vaginal bacteria (Table S4) using primer pairs (Table S3). The resulting gene products were assembled into pET28a expression vector (Novagen, 69864) using Gibson assembly (New England Biolabs, E2611S) and transformed into DH5α *E. coli* chemically competent cells. The identities of the constructs were confirmed by DNA sequencing (Genewiz). The confirmed plasmids were transformed into *E. coli* BL21(DE3) for expression. All constructs were grown in LB (Research Products International, RPI, L24060-2000) containing 50 µg/mL kanamycin sulfate (VWR, 408-EU-25G).

### Sialidases used for expression

The protein accession numbers in the NCBI database for each enzyme characterized are as follows: *G. vaginalis* NanH3 (EPI58250.1), *G. vaginalis* NanH1 (WP_013399386.1), *P. bivia* NanH (KGF23106.1), *P. denticola* (WP_036854057.1), *P. timonensi*s NanH1 (WP_008123113.1) NanH2 (WP_008122213.1).

### Heterologous Expression and Purification of Heterologously Expressed Enzymes

Chemically competent *E. coli* BL21(DE3) cells were transformed with pET28 plasmids encoding Sialidases as N-terminal His_6_-tagged fusion proteins. The transformed cells were plated onto LB_KAN_ agar plates, incubated at 37 °C overnight, and a single colony was used to grow a starter 3 mL culture and used to create a 20% glycerol stock. A 2.5 mL starter culture grown to saturation overnight was used to inoculate 1 L of LB (Research Products International, RPI) containing 50 µg/mL kanamycin. The 1 L culture was grown at 37 °C with shaking (180 rpm). When the culture reached an OD_600_ between 0.4-0.8, protein expression was induced by adding Isopropyl β-D-thiogalactoside (IPTG) (TEKNOVA, I3325) to a final concentration of 250 µM. The cultures were incubated at 16 °C with shaking (180 rpm) for 18 h. Cell cultures were then harvested by centrifugation at 6000 x g for 10 minutes at 4 °C and stored at −80 °C. Thawed cell pellets were resuspended in Buffer A containing 20 mM Tris, 500 mM NaCl, 10 mM MgSO_4_, 1 mM CaCl_2_, and 5 mM imidazole, and pH balanced at pH 7.5. The resuspended cells were supplemented with 0.1 mg/mL DNAse (Sigma, DN25-1G), 0.5 mg/mL Lysozyme (Sigma, L6876-10G), and Pierce Protease Inhibitor Tablets (Thermo Scientific, A32965). Cells were lysed using a cell disrupter (Emulsiflex-C3, Avestin) by passing three times at 15,000 psi, and cell lysates were clarified by centrifugation at 9,000 rpm for 45 minutes at 4 °C. The clarified lysate was transferred to a 15 mL column containing 3 mL Ni-NTA Agarose affinity resin equilibrated with buffer A at 4 °C (Invitrogen). The column was washed using a gradient of Buffer A containing 20 mM, 50 mM, 75 mM, and 100 mM imidazole. Protein was eluted with buffer A containing 250 mM imidazole. Fractions were analyzed by SDS-PAGE (Biorad, 4561086) and stained with InstantBlue Coomassie protein stain (Abcam, ab119211). The protein-containing fractions were pooled and concentrated in an Amicon spin concentrator of 10-30kDa cutoff (Millipore, UCF901024 and UCF903024) and buffer exchanged against Buffer B (20 mM Tris, 100 mM NaCl, 10 mM MgSO_4_, 1 mM CaCl_2_, 10 % glycerol, pH 7.5). Protein aliquots of 20 µL were frozen in liquid nitrogen and stored at –80 °C. Protein concentrations were estimated with a using NanoDrop 2000 UV-Vis Spectrophotometer (Thermo Scientific) using the theoretical molar absorption coefficient calculated using https://web.expasy.org/protparam/.

### Sialidase Activity Screen in Vaginal Isolates

Strains were grown anaerobically inside an anaerobic chamber with an atmosphere of 2.5 % H_2_, 5 % CO_2_, 92.5 % N_2_ (Coy Lab Products) in 96-well plates containing 200 µL of PYGT media supplemented with 10 % horse serum broth for 48 hours. Cultures were then normalized to OD=1 and 20 µL was added to a black 96-well plate into containing 350 µM 4-MU-Neu5Ac (Biosynth) dissolved in 80 µL sodium acetate buffer pH 5.5. Fluorescence signal was monitored at 360/440 nm for 105 minutes in a plate reader (Biotek Synergy HTX) at 37 °C.

### Activity Assays of Purified Sialidases

Sialidase activity was measured using a fluorescence-based assay in a total volume of 100 µL, prepared in a black, 96-well polystyrene plate (Corning). Each well contained 2.5 nM purified enzyme in sodium acetate buffer (100 µM, pH 5.5) and 4-MU-Neu5Ac (200 µM) was added to start the sialidase reaction and fluorescence 330/440 nm was measured while incubating at 37 °C with shaking in a plate reader (Biotek).

### Kinetic Characterization

Sialidase activity was measured using 2.5 nM purified enzyme in sodium acetate buffer (100 µM, pH 5.5) to a total volume of 100 µL. 4-MU-Neu5Ac was diluted to create a 12-step, 1:2 dilution series from 800 µM to 3.1 µM. Fluorescence 360/440 nm was measured while incubating at 37 °C with shaking in a plate reader (Biotek). Slopes were calculated over the first 5 minutes and used GraphPad to calculate Michaelis–Menten kinetic parameters, Vmax and Km.

### pH Profiles of Sialidases

The optimal pH for enzyme activity was determined by incubating 20–100 nM sialidases with 200 µM 4-MU-Neu5Ac in Mcllvaine buffer (0.1 M citric acid / 0.2 M phosphate buffer) at a pH range of 3 to 8 for 30 minutes in at 37 °C. Activity was assayed by monitoring fluorescence 360/440 nm in a plate reader (Biotek). The assay was prepared in a 384-well, black, flat bottom polystyrene plate (Corning). Fluorescence values were normalized to standard curves of 4-methylumbelliferone (Sigma) (concentration 200 µM to 1.5 µM) dissolved in DMSO and prepared in the same citrate/phosphate buffers from pH 3 to 8. Relative sialidase activity was determined by measuring the total concentration of 4-methylumelliferone released at 29 min.

### Sialidase Substrate Specificity Endpoint Assay

To determine the substrate specificity of sialidases, 40 nM enzyme was incubated with the following substrates: 4-MU-Neu5Ac (250 µM), BSM (1 mg/mL), human serum IgA (0.5 mg/mL), human serum IgG (4 mg/mL), 3’sialyllactose (200 µM), 6’siallylactose (200 µM), MUC5B (0.5 mg/mL) in 20 mM sodium acetate buffer pH 5.5 in a total volume of 50 µL. All samples included *N*-Acetyl-D-neuraminic acid-1,2,3-^13^C_3_ (Sigma, 649694) as an internal standard. The assays were incubated for 2 hours at 37 °C in a thermocycler (Biorad). Samples were then filtered through 3 kDa filter plates and derivatized with 4,5-dimethoxy-1,2-phenylenediamine hydrochloride (DMB), (Sigma, 36271). To derivatize samples, 20 µL of filtered sample was mixed with 80 µL of derivatization reagent (6.5 mM DMB, 52 mM sodium dithionite, 0.75 M 2-mercaptoethanol, 1.33 M acetic acid). Samples were incubated at 50 °C for 2 hours.

### Anaerobic Culture Conditions

All plastic consumables used for culturing were kept in the anaerobic chamber for at least 24 hours prior to the experiment. Bacterial isolates were obtained from Biodefense and Emerging Infections Research Resources Repository (BEI Resources), (Table S3). *Gardnerella* and *Prevotella* strains were cultured in PYGT media as previously described^40^ (20 g peptone, 10 g glucose, 10 g yeast extract, 0.4 g sodium bicarbonate, 40 mg dipotassium phosphate, 40 mg monopotassium phosphate, 5 mg hemin, 5 mg vitamin K, 0.25 g ʟ-cysteine hydrochloride, 8 mg magnesium sulfate, 0.250 mL tween-80, 50 mL heat inactivated horse serum (Sigma), in 1 L water filter sterilized (0.2 µm), containing 10 % inactivated horse serum. Media was transferred to an anaerobic chamber with an atmosphere containing 2.5 % H_2_, 5 % CO_2_ and 92.5 % N_2_ (Coy Labs). Starter cultures of *Gardnerella* and *Prevotella* were inoculated in 96-well plates containing 200 µL of PYGT and incubated at 37°C overnight.

### Sialic Acid Detection by DMB labeling and LC-MS analysis

Derivatized samples were prepared for analysis by diluting 1:100 in 90:10 acetonitrile (ACN): water. Samples were analyzed by ultra-high performance model Xevo TQ-S (Waters, UPLC-MS/MS) see supplemental methods and materials (Table S4).

### MUC5B purification

Submandibular saliva was collected in bulk from human volunteers using a custom vacuum pump, as described previously^41,42^. Immediately after collection, salts, antibacterial agents, and protease inhibitors were added to the saliva to reach a final concentration of 0.16 M NaCl, 5 mM benzamidine HCl, 1 mM dibromoacetophenone, 1 mM phenylmethylsulfonyl fluoride, and 5 mM EDTA. The mucins in the saliva were solubilized overnight by stirring gently at 4 °C. Solubilized saliva was then flash-cooled in liquid nitrogen in 10-40 mL volumes and stored at -80°C. Before chromatography, 200 mL of saliva from separate donors was thawed at 4°C, and insoluble material was removed by centrifugation at 10,000 × g for 10 min at 4 °C. MUC5B was purified using a Bio-Rad NGC fast protein liquid chromatography (FPLC) system equipped with an XK 50 column packed with 2 L of Sepharose CL-2B resin (GE Healthcare Bio-Sciences). Mucin-containing fractions were identified using a periodic acid-Schiff’s reagent assay and analysis of UV absorbance at 280 nm from FPLC. Fractions were then combined, dialyzed, and concentrated using an ultrafiltration device and were then lyophilized for storage at −80°C. Protocols involving the use of human subjects were approved by Massachusetts Institute of Technology’s Committee on the Use of Humans as Experimental Subjects.

### Inhibition of Purified Sialidases

Sialidase inhibition assays were performed using 5 nM purified enzyme in sodium acetate buffer (100 µM, pH 5.5) and with varying concentrations of Neu5ac2en (Sigma, D9050) in a total volume of 50 µL. Each well contained 1 µL of enzyme was pre-mixed with 39 µL of buffer and 5 µL of inhibitor in a black 384-well flat bottom polystyrene plate (Corning) and 5 µL of 4-MU-Neu5Ac (100 µM final concentration) was added to start the assay. The plate was immediately transferred into the plate reader to incubate at 37 °C with shaking to measure fluorescence 360/440 nm. For improved accuracy, the enzyme, inhibitor, and 4-MU-Neu5Ac were dispensed into the plate using a Formulatrix MANTIS. Slopes to determine sialidase activity were calculated over the first 5 minutes and the IC_50_ were determined by non-linear fit [Inhibitor] vs. response on Graphpad Prism.

### Bacterial genomic database and HMM-based sialidase searches of isolate genomes

A bacterial isolate genomic database was constructed using 1,189 published genomes from the Vaginal Microbiome Research Consortia, spanning cohorts from the United States and South Africa (Full list and metadata provided in Supplementary data file). Bacterial genomes were organized by phylogeny (Figure S9-10) using a concatenated ribosomal protein tree. We used HMMER (v3.3.2) to find ribosomal proteins, aligned the sequences with MAFFT (v7.508) and used RAxML (v.8.2.10) to create the phylogenetic tree in Figure S9-10. In order to search our diverse collection of vaginal bacterial genomes or putative sialidases, we used HMMER (v3.3.2) to construct a hidden Markov model using a database made of a multiple sequence alignment of nine sialidase genes (See supplementary fasta file A) constructed with MAFFT (v7.508). We required all hits to be greater than 250 amino acids in length (See supplementary fasta file B) and validated hits manually by searching for predicted sialidase domains using InterProScan.

### Metagenome and Metatranscriptome Searches and Quantification

Diamond blastX was used to quantify the abundance of sialidase encoding genes and transcripts in metagenomes and metatranscriptomes published by the Vaginal Microbiome Research Consortium (VMRC). First, we compiled all the candidate sialidase genes identified by HMMER and created representative sequences using CD-hit with >85 % amino acid ID. After representative sequences were generated, they were used in Diamond blastX with a e-value cutoff of <E-20 and a percent amino acid sequence identity >50 % to determine the abundance of these sialidases in metagenome and metatranscriptomes databases generated by the VMRC under the Bioproject PRJNA797778. By setting a stringent identity and e value cutoff, we are quantifying high confidence hits to specific sialidase genes queried. The scripts for processing the datasets were described previously^19^. We analyzed 176 paired metagenomes and metatranscriptomes from 40 patients. The output from diamond blastx is RPKM. Sample metadata was used to bin the results by community state type (CSTI n=39, CSTII n=16, CSTIII n=31, CSTIV n=80, CSTV n=10). The total expression per sample was calculated by summing all the RPKM values for all the candidate sialidases for each sample. Percent contribution of a sialidase is determined by the relative abundance of one sialidase divided by the sum of all sialidase abundances within each sample. Plots were generated using Python 3 and PRISM.

## Acknowledgments

We thank Karin Strijbis and Celia Sequi Perez for critical reading and editing of the manuscript. We thank Wei Li for assistance with cloning the *Gardnerella* sialidases. We acknowledge Beverly Fu, Min Woo Bae, and Grace Kenney for providing advice regarding bioinformatic analyses. We thank all the members of the Vaginal Microbiome Research Consortium for helpful discussions. We thank Michael France and Jacques Ravel for sharing the metagenomic and metatranscriptomic datasets and metadata. We acknowledge funding from the Bill and Melinda Gates Foundation (awards No. 270790 and INV-037720). E.P.B. is a Howard Hughes Medical Institute Investigator. P.P. was supported by the National Science Foundation Graduate Research Fellowship (NSF-GRFP). F.A.H was supported by the Schmidt Science Fellowship. D.S.K. was supported by the Bill and Melinda Gates Foundation (awards INV-048977 and OPP1189208). This article is subject to HHMI’s Open Access to Publications policy. HHMI lab heads have previously granted a nonexclusive CC BY 4.0 license to the public and a sublicensable license to HHMI in their research articles. Pursuant to those licenses, the author-accepted manuscript of this article can be made freely available under a CC BY 4.0 license immediately upon publication.

