## Supplementary Material for "*Prevotella* are major contributors of sialidases in the human vaginal microbiome"

### Supporting Information for
*Prevotella* are major contributors of sialidases in the human vaginal microbiome.

Paula Pelayo[1], Fatima A. Hussain[2], Caroline A. Werlang[3], Chloe Wu[3], Benjamin M. Woolston[1,4], Katharina Ribbeck[3], Douglas S. Kwon[2], Emily P. Balskus[1,5].

Emily P. Balskus.


**This PDF file includes:**

Supplemental Methods and Materials
Figures S1 to S12
Tables S1 to S8
SI References

**Other supporting materials for this manuscript include the following:**

Supplementary data file.xls
Supplementary fasta file A.faa
Supplementary fasta file B.faa

#### Supplemental Methods and Materials

All chemicals and solvents were purchased from Sigma-Aldrich, except where otherwise noted.

**Table S1.** Primers used for cloning sialidases.

| Name | Description | Sequence (5' to 3') |
|---|---|---|
| oPP_01 | pET28a-F (N-His + Thrombin) | GCTGCCGCGCGGCACCAG |
| oPP_02 | pET28a-R (Eliminate other tags) | CAAAGCCCGAAAGGAAGCTG |
| oPP_03 | P. bivia DNF00188 sialidase-F | cacagcagcggcctggtgccgcgcggcagcCAAGATACCTTGTTTGTTAAAC |
| oPP_04 | P. bivia DNF00188 sialidase-R | gcagccaactcagcttcctttcgggctttgTTATTTCTCTTTGATAATATCT |
| oPP_05 | P. denticola DNF00960 sialidase-F | cacagcagcggcctggtgccgcgcggcagcGCCGGGACAGACAGGAACAGAG |
| oPP_06 | P. denticola DNF00960 sialidase-R | gcagccaactcagcttcctttcgggctttgTCAGCGGCGACGTGCGGCAGCG |
| oPP_07 | P. timonensis cris 5c-b1 sialidase long-F | cacagcagcggcctggtgccgcgcggcagcGCGGACAAGGTAATCCGCATTC |
| oPP_08 | P. timonensis cris 5c-b1 sialidase long-R | gcagccaactcagcttcctttcgggctttgTTACTTCACCACCACCTTTTTG |
| oPP_09 | P. timonensis cris 5c-b1 sialidase short-F | cacagcagcggcctggtgccgcgcggcagcTCAAACAACCAGCATCACCAAC |
| oPP_10 | P. timonensis cris 5c-b1 sialidase short-R | gcagccaactcagcttcctttcgggctttgTTAGTAGCCCTTCTTGAAGCGT |
| oBMW_011 | G. vaginalis JCP8066 NanH3-F | cagcggcctggtgccgcgcggcagcACTACCCCCCCCATGAAC |
| oBMW_012 | G. vaginalis JCP8066 NanH3-R | caactcagcttcctttcgggctttgTTAATATTTCATATTTTTTAATTTCATTAATGCTGTAGAGATAGCTATTAATAGAAG |
| oBMW_013 | G. vaginalis ATCC14018 NanH1-F | cagcggcctggtgccgcgcggcagcATGGAACGTCGTTCAACG |
| oBMW_014 | G. vaginalis ATCC14018 NanH1-R | caactcagcttcctttcgggctttgTTAAATGTCTCTTCCATGTTGG |

**Table S2.** Bacterial strains used for sialidase characterization and assays of sialidase activity.

| Sialidase | Cloned from this strain | GenBank assembly Accession | Source |
|---|---|---|---|
| *Gv*NanH3 | *G. vaginalis JCP8066* | GCF_000414565.1 | Biodefense and Emerging Infections Research Resources Repository (BEI Resources) |
| *Gv*NanH1 | *G. vaginalis ATCC14018* | GCA_003397685.1 | BEI Resources |
| *Pb*NanH | *P. bivia DNF00188* | GCA_000759045.1 | BEI Resources |
| *Pd*NanH | *P. denticola DNF00960* | GCA_000759205.1 | BEI Resources |
| *Pt*NanH1 | *P. timonensis cris 5c-b1* | GCA_000177055.1 | BEI Resources |
| *Pt*NanH2 | *P. timonensis cris 5c-b1* | GCA_000177055.1 | BEI Resources |

**Table S3.** Cloning strains (A) and (B) plasmids used in this study.

A. *E. coli* strains

| Name | Description / Use | Source |
|---|---|---|
| *E. coli* DH5α | Cloning strain | NEB (New England BioLabs) |
| *E. coli* BLL21(DE3) | Protein expression and purification | NEB |

B. Plasmids

| Plasmid | Description | Source |
|---|---|---|
| pET28a-*GvnanH1* | N-$His_6$, thrombin, *G. vaginalis nanH1*, $Km^r$ | This study |
| pET28a-*GvnanH3* | N-$His_6$, thrombin, *G. vaginalis nanH3*, $Km^r$ | This study |
| pET28a-*PbnanH* | N-$His_6$, thrombin, *P. bivia nanH*, $Km^r$ | This study |
| pET28a-*PdnanH* | N-$His_6$, thrombin, *P. denticola nanH*, $Km^r$ | This study |
| pET28a-*PtnanH1* | N-$His_6$, thrombin, *P. timonensis nanH1*, $Km^r$ | This study |
| pET28a-*PtnanH2* | N-$His_6$, thrombin, *P. timonensis nanH2*, $Km^r$ | This study |

**UPLC-MS/MS methods for detecting derivatized sialic acids**

Derivatized samples were prepared for analysis by diluting 1:100 in 90:10 acetonitrile (ACN): water. Samples were analyzed by ultra-high performance liquid chromatography tandem mass spectrometry (UPLC-MS/MS). Simultaneous analysis of Neu5Ac, *N*-Acetyl-D-neuraminic acid-1,2,3-$^{13}C_3$ (Sigma, 649694), and Neu5Gc derivatized with 4,5-dimethoxy-1,2-phenylenediamine hydrochloride (DMB) was carried out by UPLC-MS/MS. Liquid chromatography was conducted using a Waters Aquity H-Class System (Waters Corporation). Following sample preparation, 1 µL of sample was injected onto a BEH Amide 1.7 µm (2.1 x 50 mm) column (Acquity, 186004801). The flow rate was 0.650 mL $min^{-1}$ using mobile phase A = 0.1 % formic acid in $H_2O$ and mobile phase B = 0.1 % formic acid in acetonitrile (ACN). The column temperature was maintained at 40 °C. The following gradient was applied: 0–1.5 min at 90-60% B, 1.5-2 min at 60% B isocratic, 2.0-2.2 min at 60-90% B, 2.2–3.5 min at 90% B isocratic. MS detection was performed with a Waters Xevo TQ-S (Waters Corporation) instrument with electron spray ionization in positive mode (ESI+) (capillary voltage,3.3 kV; cone voltage, 48 V; source offset voltage, 50 V; desolvation temperature, 450 °C; desolvation gas flow, 250 L h–1; cone gas flow, 150 L h–1; nebulizer, 4.0 bar). For quantification, standard curves for Neu5Ac and Neu5Gc ranging from 0.1–200 µM were prepared in triplicate. Standards were prepared and derivatized in parallel with experimental samples for each experiment.

**Table S4.** UPLC-MS/MS analysis of sialic acids used in assays.

| *Standard* | *Transition (m/z)* | *Mode* | *Cone (V)* | *Collision (V)* | *Retention (min)* |
|---|---|---|---|---|---|
| DMB-Neu5Ac | 442.33>424.298 | ESI | 48 V | 10 | 1.57 |
| DMB-$^{13}$CNeu5Ac | 445.33>427.298 | ESI | 48 V | 10 | 1.55 |
| DMB-Neu5Gc | 458.17> 440.17 | ESI | 48 V | 10 | 1.67 |

**Inhibition of sialidase activity in bacterial culture**

Bacteria were grown in PYGT media for 48 hours, diluted to an OD=1 and then 10 µL of culture was added to a black 384-well flat bottom polystyrene plate (Corning) containing sodium acetate buffer (100 µM, pH 5.5) and with varying concentrations of Neu5ac2en (Sigma, D9050) in a total volume of 50 µL. To start the assay, 5 µL of 4-MU-Neu5Ac (100 µM final concentration) was added and the plate was immediately transferred into the plate reader to incubate at 37 °C with shaking to measure fluorescence 360/440 nm for 2 hours. For improved accuracy, the 4-MU-Neu5Ac substrate was dispensed into the plate using a Formulatrix MANTIS. Slopes to determine sialidase activity were calculated over 20 points across the 2 hours and the $EC_{50}$ were determined by non-linear fit [Inhibitor] vs. response on Graphpad Prism.

**Supplemental Figures**

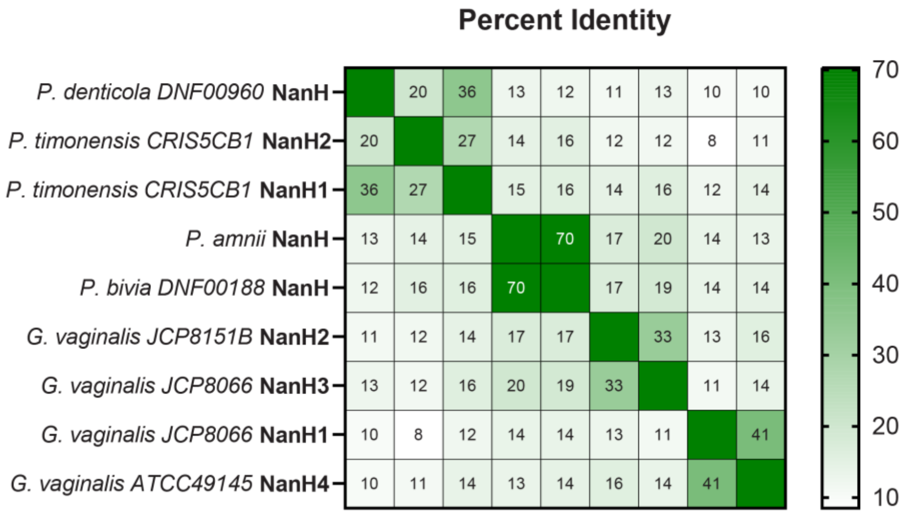


**Figure S1**. Muscle alignment of the full amino acid sequences from 4 *Gardnerella* and 5 *Prevotella* sialidases.

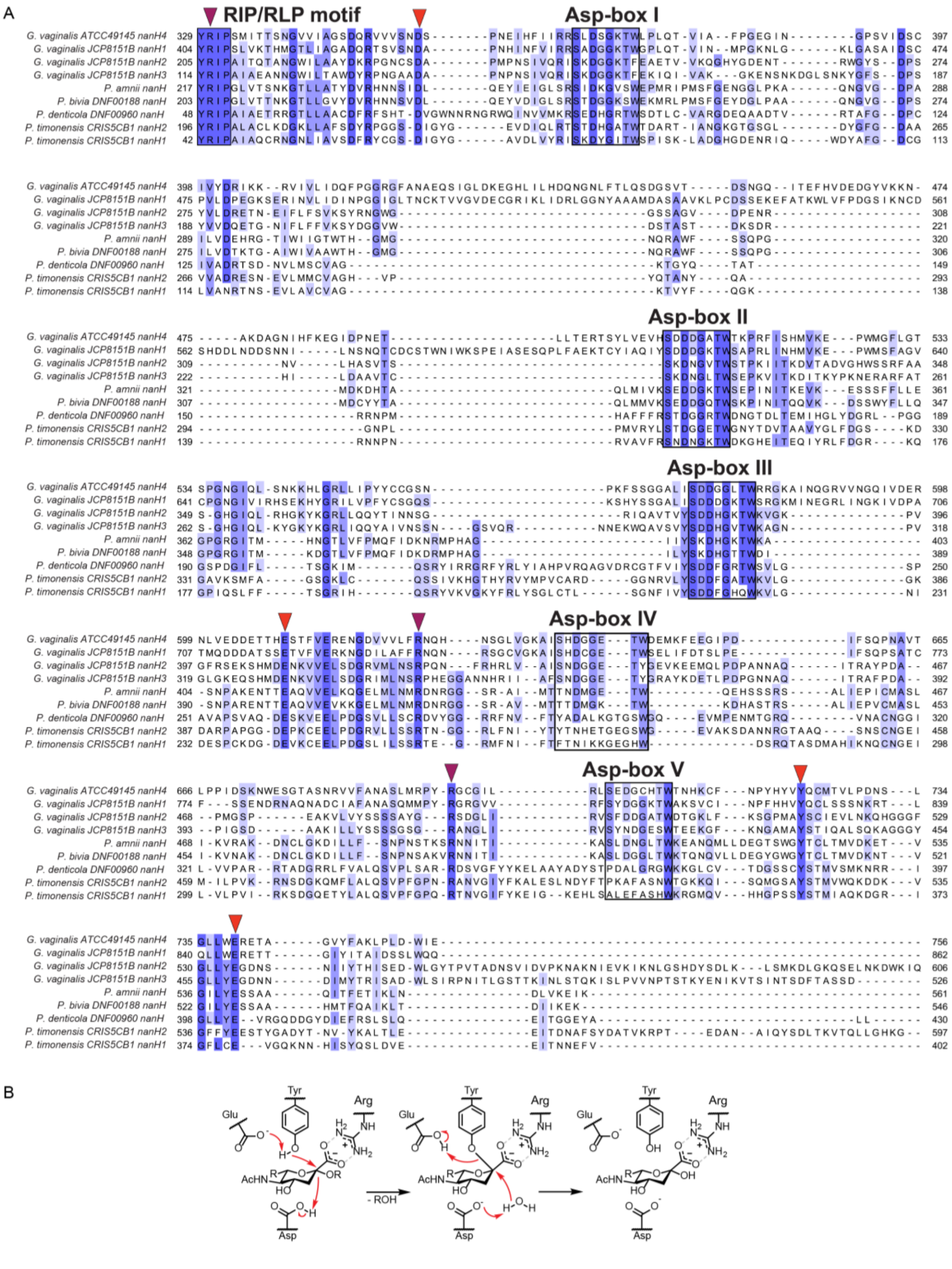


**Figure S2.** Multiple sequence alignment of 4 *Gardnerella* and 5 *Prevotella* sialidases showing the conserved active site residues involved in catalysis (red and maroon arrows). **A)** Rectangles indicate the conserved sequence motifs, RIP/RLP motif (Arg-Ile/Leu-Pro) and the signature bacterial neuraminidase repeat, BNR repeat (Ser/Thr-X-Asp-X-Gly-X-Thr-Trp/Phe). BNR repeats are also known as Asp-box motifs[1]. **B)** Proposed mechanism for sialidases.

**Table S5.** Active site residues conserved among *Gardnerella* and *Prevotella* sialidases.

| Organism | Sialidase | Arg triad | Tyr/Glu-E | Asp-D | Glu |
|---|---|---|---|---|---|
| *G. vaginalis* | NanH1 | 405/733/800 | 828/717 | 430 | 844 |
| *G. vaginalis* | NanH2 | 206/423/487 | 515/407 | 231 | 534 |
| *G. vaginalis* | NanH3 | 115/345/412 | 440/329 | 140 | 459 |
| *G. vaginalis* | NanH4 | 330/695/ | 723/609 | 355 | 732 |
| *P. bivia* | NanH | 204/415/479 | 510/399 | 229 | 526 |
| *P. timonensis* | NanH1 | 43/257/325 | 362/241 | 67 | 378 |
| *P. timonensis* | NanH2 | 197/412/485 | 524/396 | 221 | 540 |
| *P. denticola* | NanH | 49/276/347 | 386/260 | 73 | 402 |

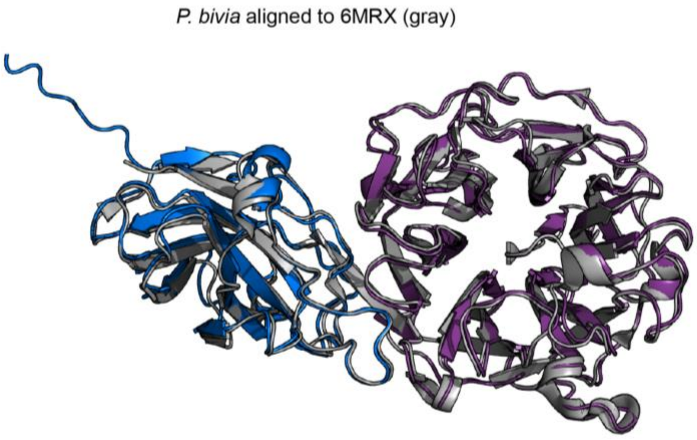


**Figure S3.** Predicted structures of *P. bivia* sialidase PbNanH generated via ColabFold AlphaFold2. **A)** Sialidase domain IPR01140 (purple) is present, forming the catalytic β-propeller fold domain. PbNanH contains an additional predicted sialidase carbohydrate binding domain (blue). PbNanH shares 66% amino acid ID with Sialidase26 (6MRX), aligned in gray[2].

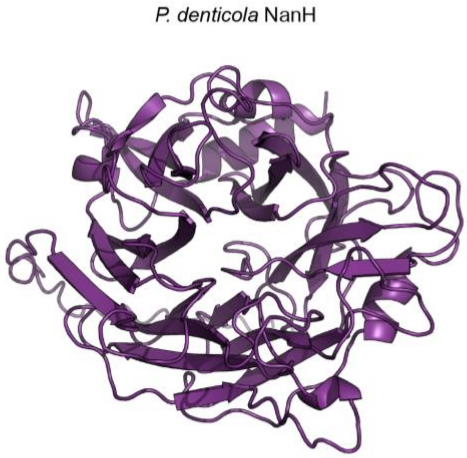


**Figure S4.** Predicted structures of *P. denticola* sialidase PdNanH generated via ColabFold AlphaFold2. Sialidase domain IPR01140 is present, forming the catalytic β-propeller fold domain.

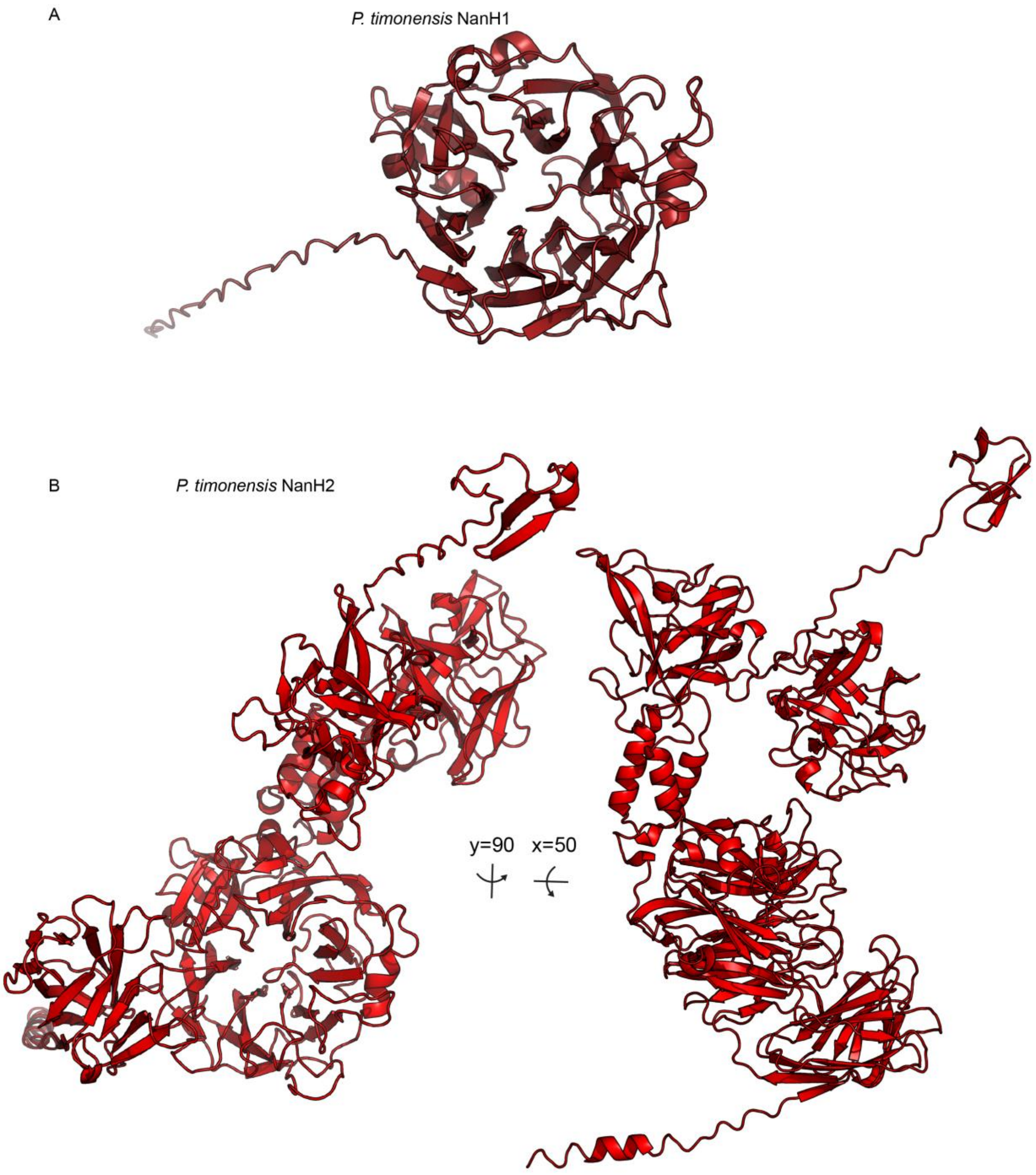


**Figure S5.** Predicted structures of *P. timonensis* sialidases generated via ColabFold AlphaFold2 **A)** PtNanH1 and **B)** PtNanH2. PtNanH2 contains the IPR01140 domain and 4 additional structural elements of unknown function.

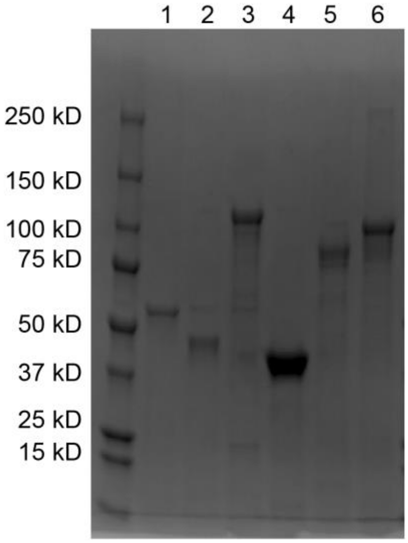


**Figure S6**. SDS-PAGE of recombinant purified sialidases. From left to right, Precision Plus Protein All Blue Standards (BioRad), (1) *P. bivia* NanH, 60.7 kD; (2) *P. denticola* NanH, 49 kD; (3) *P. timonensis* NanH2, 111 kD; (4) *P. timonensis* NanH1, 46 kD; (5) *G. vaginalis* NanH3, 79 kD; (6) *G. vaginalis* NanH1, 101 kD. Gel was stained with Instant Blue (Abcam).

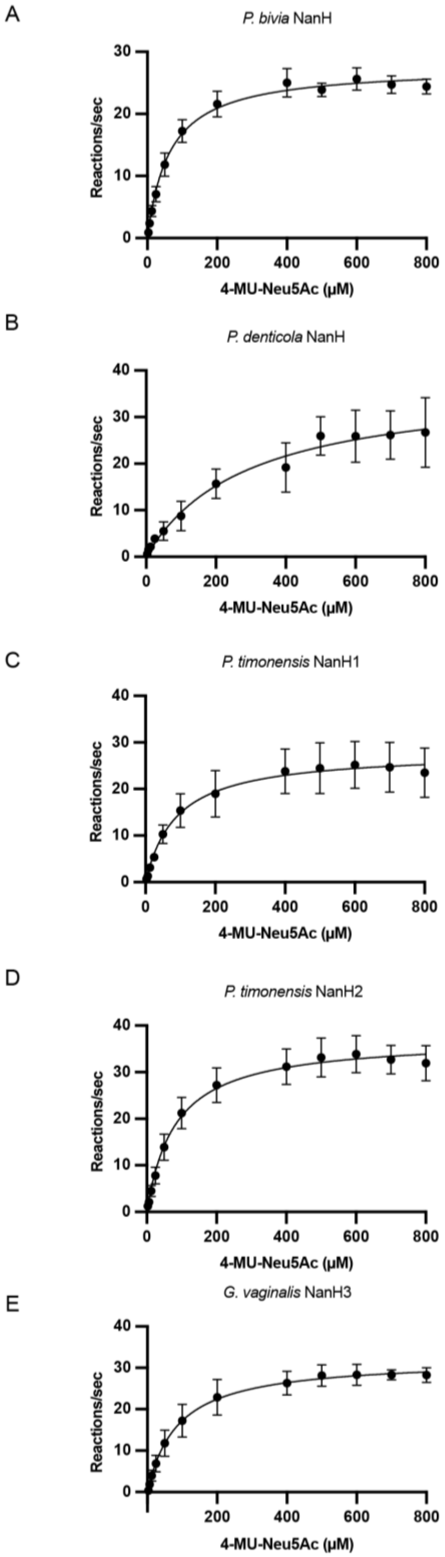


**Figure S7.** Michaelis–Menten kinetic parameters for sialidases. A) Kinetic characterization of purified sialidases. 2.5 nM of each enzyme was incubated with varying concentrations of 4-MU-Neu5Ac for 5 minutes at 37 °C. Mean values are shown as filled circles +/- standard error of means (SEM) from three independent experiments (n=3) using the same batch of purified enzymes on the same day.

**Table S6.** Kinetic parameters of sialidases for the hydrolysis of 4-MU-Neu5Ac. Values represent mean of three independent experiments on the same day (n=3). Range represents ± SD.

| Enzyme | $k_{cat}$ ($s^{-1}$) | Catalytic efficiency $k_{cat}$ / $K_m$ ($s^{-1}$ $M^{-1}$) | $K_m$ (μM) | $V_{max}$ (μM $min^{-1}$) |
|---|---|---|---|---|
| *P. timonensis* NanH1 | 112.20 ± 8.29 | (1.26 ± 0.24) x $10^6$ | 88.90 ± 27.14 | 16.83 ± 1.24 |
| *P. timonensis* NanH2 | 149.20 ± 6.28 | (1.83 ± 0.40) x $10^6$ | 81.37 ± 14.53 | 22.38 ± 0.94 |
| *P. bivia* NanH | 110.46 ± 2.87 | (1.73 ± 0.25) x $10^6$ | 64.01 ± 7.48 | 16.57 ± 0.43 |
| *P. denticola* NanH | 149.33 ± 22.21 | (0.51 ± 0.28) x $10^6$ | 290.30 ± 113.43 | 22.40 ± 3.33 |
| *G. vaginalis* NanH3 | 128.60 ± 5.38 | (1.48 ± 0.32) x $10^6$ | 86.79 ± 15.10 | 19.29 ± 0.81 |

**Table S7.** For comparison, kinetic parameters from previously characterized sialidases for the hydrolysis of *p*-nitrophenyl-*N*-acetylneuraminic acid (*p*-NP-Neu5Ac).

| Organism | Enzyme | $k_{cat}$ ($s^{-1}$) | Catalytic efficiency $k_{cat}$ / $K_m$ ($s^{-1}$ $M^{-1}$) | $K_m$ (μM) | Substrate | Study |
|---|---|---|---|---|---|---|
| *Streptoccoccus pneumoniae* | NanA | >175 | (3.5 ± 0.3) x $10^5$ | >500 | *p*-NP-Neu5Ac | 3 |
| *Streptoccoccus pneumoniae* | NanB | >0.14 | (2.7 ± 0.3) x $10^2$ | >500 | *p*-NP-Neu5Ac | 3 |
| *Streptoccoccus pneumoniae* | NanC | >17 | (3.4 ± 0.3) x $10^4$ | >500 | *p*-NP-Neu5Ac | 3 |

**Table S8.** Activity of sialidase inhibitors (up to 2 μM) towards the activity of purified vaginal sialidases on 4-MU-Neu5Ac. ND = Not determined.

| | $IC_{50}$ (μM) | |
|---|---|---|
| | Neu5Ac2en | Zanamivir |
| *P. bivia* NanH | 103.3 ± 20.3 | ND |
| *P. denticola* NanH | 50.5 ± 22.5 | ND |
| *P. timonensis* NanH1 | 180.0 ± 92.3 | 4426.0 ± ND |
| *P. timonensis* NanH2 | 56.0 ± 23.0 | 33.6 ± 15.5 |
| *G. vaginalis* NanH3 | 29.0 ± 5.6 | ND |

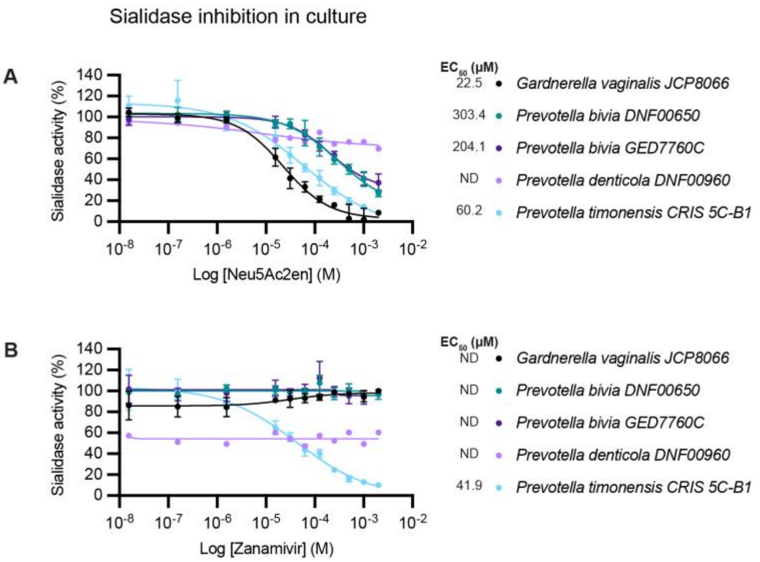


**Figure S8.** Inhibitory activity of (A) Neu5Ac2en and (B) Zanamivir towards *Prevotella* and *Gardnerella* cultures.

Bacteria grown in PYGT media for 48 hours were incubated with varying concentrations of Neu5ac2en and Zanamivir for 2 h at 37 °C. Data represent the mean of 3 biological replicates ± SEM (except *P. denticola* which only has one replicate).

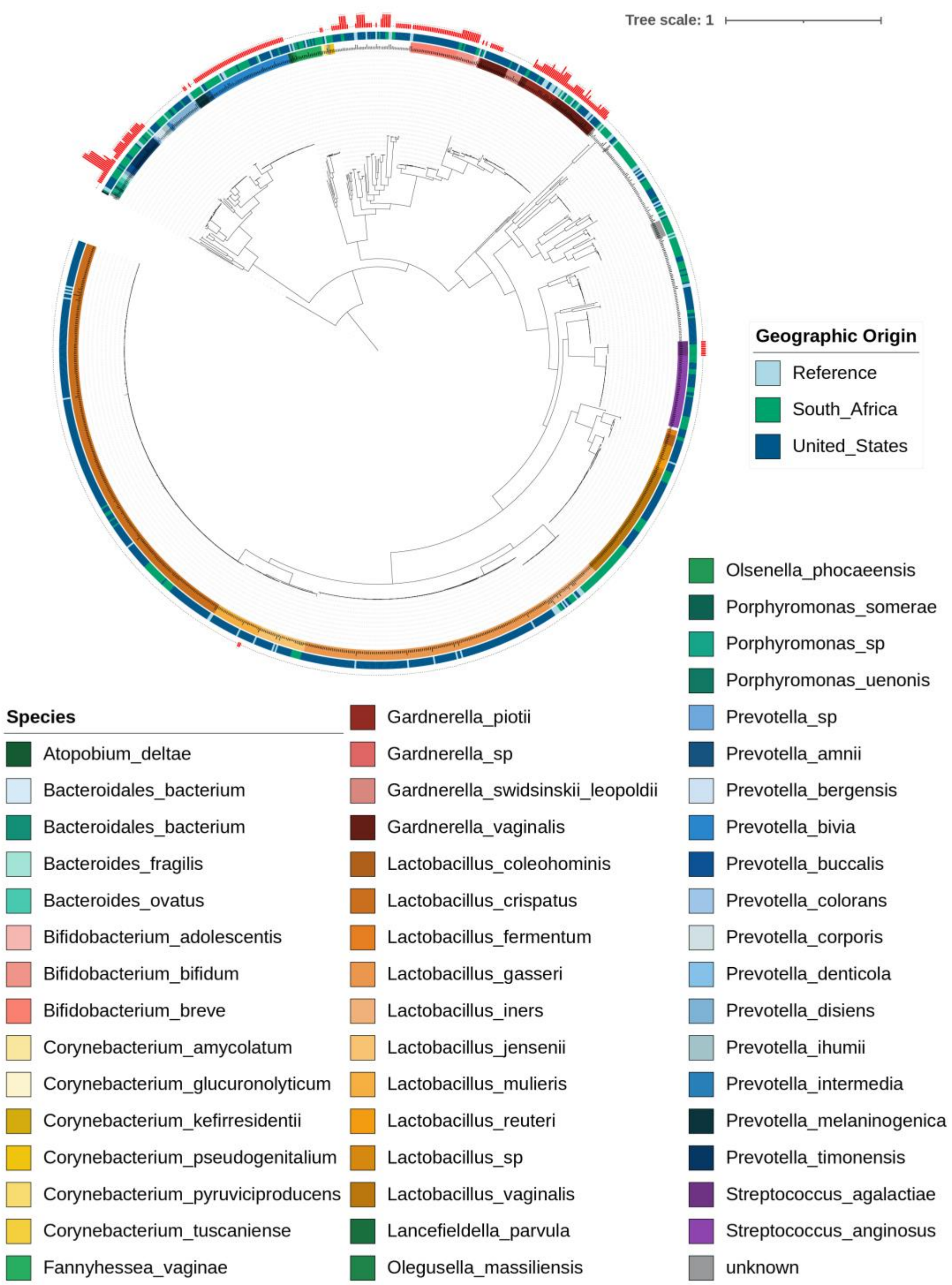


**Figure S9.** Sialidases encoded in vaginal isolate genomes.
Phylogenetic tree of vaginal isolates from the Vaginal Microbiome Research Consortium (N. American) and FRESH (S. African) studies. The phylogeny is based on 49 concatenated ribosomal proteins and serves as a proxy for the core genome. Each genome was searched using the sialidase protein alignment with HMMER (version 3.1b2). The scale bar indicates nucleotides substitutions per site. The presence of sialidases is represented by the red bar next to the isolate ID with the height and number indicating the number of sialidase hits.

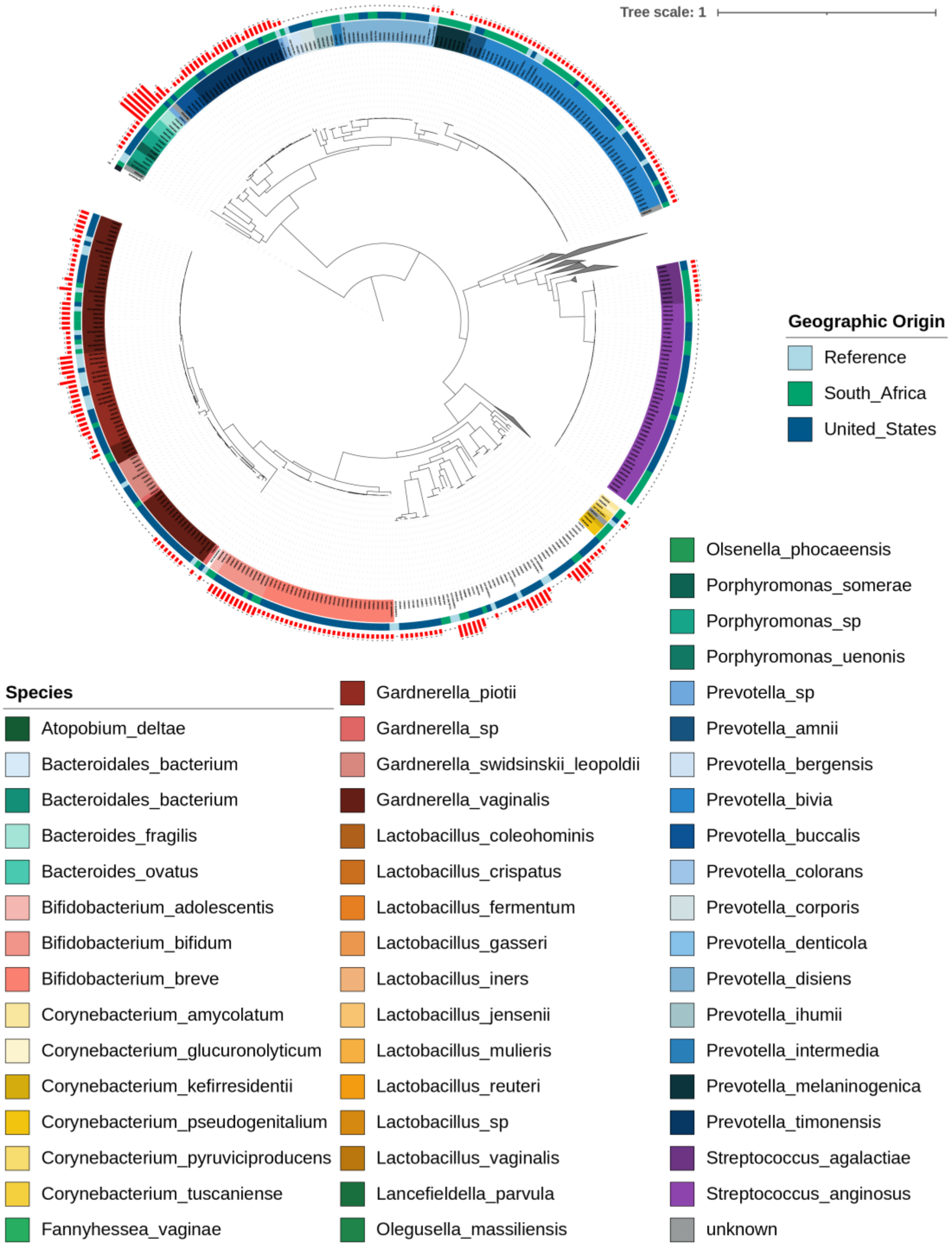


**Figure S10.** Sialidases encoded in vaginal isolate genomes with *Lactobacillus* clade collapsed
This figure is identical to Figure S8, but with the *Lactobacillus* clade collapsed. Sialidases are shown in red bars next to the isolate ID and the height indicates the number of sialidase hits. Phylogenetic tree of vaginal isolates from the Vaginal Microbiome Research Consortium (N. American) and FRESH (S. African) studies. The phylogeny is based on 49 concatenated ribosomal proteins and serves as a proxy

for the core genome. Each genome was searched using the sialidase protein alignment with HMMER (version 3.1b2). The scale bar indicates nucleotides substitutions per site.

A

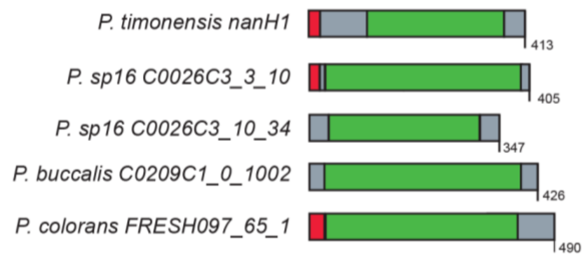


B

| MUSCLE amino acid alignment | | | | | |
|---|---|---|---|---|---|
| *P. timonensis*_nanH1 | *P. timonensis* nanH1 | *P. sp16* C0026C3_3_10 | *P. sp16* C0026C3_10_34 | *P. buccalis* C0209C1_0_1002 | *P. colorans* FRESH097_65_1 |
| *P. sp16* C0026C3_3_10 | 56.2 | | | | |
| *P. sp16* C0026C3_10_34 | 22.8 | 20.1 | | | |
| *P. buccalis* C0209C1_0_1002 | 61.6 | 54.9 | 20.2 | | |
| *P. colorans* FRESH097_65_1 | 9.5 | 10.6 | 10.6 | 9.5 | |

**Figure S11.** Predicted sialidase domains among new *Prevotella* sialidases from genome survey in Figure 4.
A) *Prevotella* encode several predicted sialidases with sialidase domains (green) and signal peptides (red). *P. timonensis* NanH1 was characterized in this study. B) MUSCLE alignment of proteins, the numbers represent % amino acid identity.

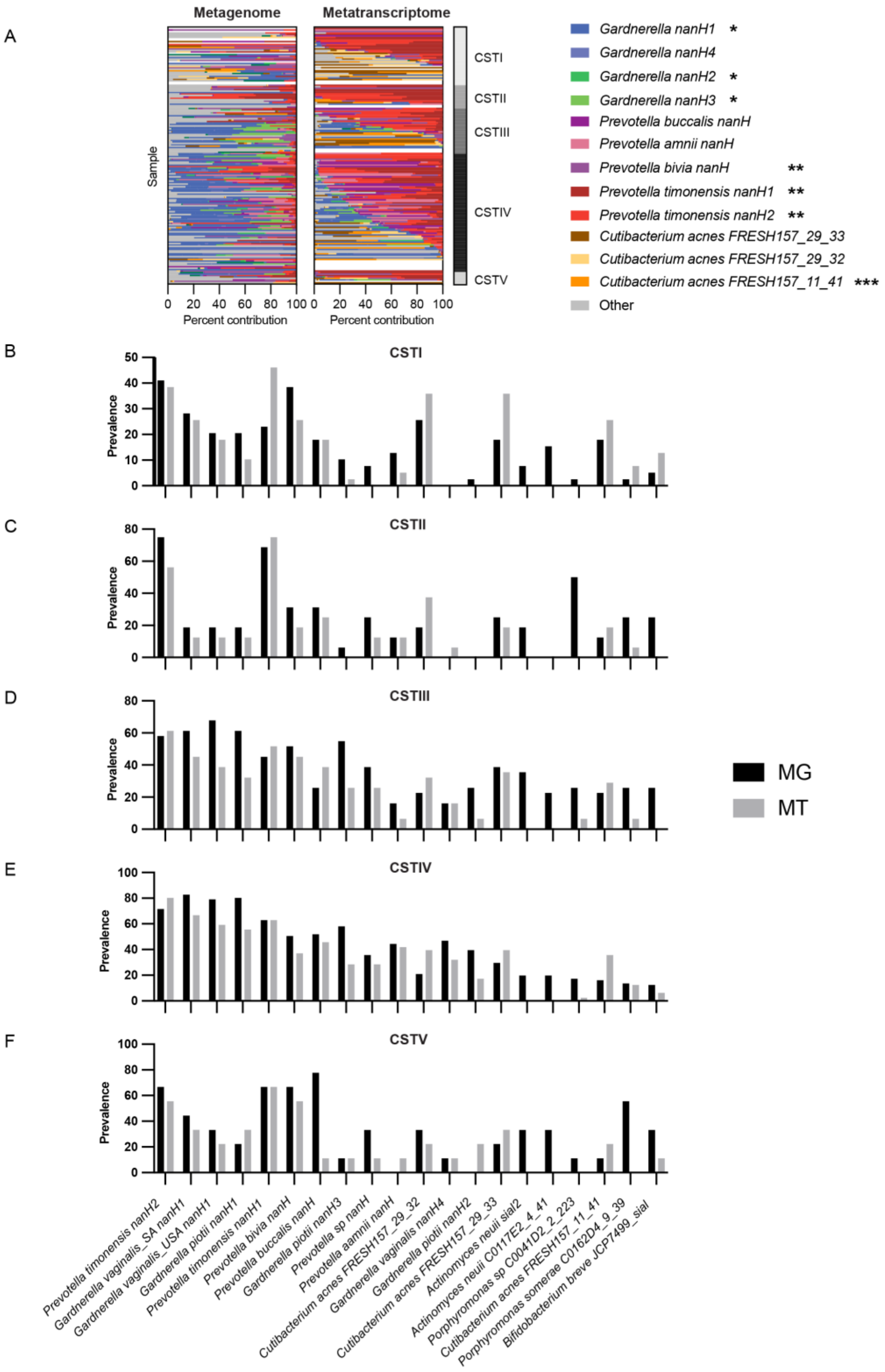


**Figure S12.** *Prevotella* sialidases are prevalent across all vaginal community state types.
(A) Contribution of specific vaginal sialidase in metagenomes (left) and metatranscriptomes (right). Paired metagenomes (MG) and metatranscriptomes (MT) n=176 were used to investigate the relative contribution of several vaginal sialidases. * indicates enzymes were characterized previously [4], **

characterized in this study, *** indicates a *Cutibacterium acnes* sialidase with 99.6% a.a. ID to formerly characterized *Propionibacterium acnes* sialidase (PaNa, GenBank: ATT83304.1)[5]. This panel displays the same data in Figure 5D, in more detail to display the resolution of specific genes present in samples. Prevalence of select sialidase genes with >15 % prevalence in MG samples found in (B) CSTI (C) CSTII (D) CSTIII (E) CSTIV (F) CSTV. This panel displays the same data in Figure 5A, binned by community state type.
